# Mechanochemical forces regulate the composition and fate of stalled nascent chains

**DOI:** 10.1101/2024.08.02.606406

**Authors:** Danish Khan, Ananya A Vinayak, Cole S Sitron, Onn Brandman

**Affiliations:** Department of Biochemistry, Stanford University School of Medicine, Stanford, CA 94305, USA; Department of Cellular Biochemistry, Max Planck Institute of Biochemistry, Am Klopferspitz 18, 82152 Martinsried, Germany

**Keywords:** CAT tails, protein quality control, ribosome, translation, ribosome stalling, mechanochemistry, ribosome-associated quality control (RQC), protein folding

## Abstract

The ribosome-associated <u>q</u>uality <u>c</u>ontrol (RQC) pathway resolves stalled ribosomes. As part of RQC, stalled nascent polypeptide chains (NCs) are appended with <u>CA</u>rboxy-<u>T</u>erminal amino acids (CAT tails) in an mRNA-free, non-canonical elongation process. CAT tail composition includes Ala, Thr, and potentially other residues. The relationship between CAT tail composition and function has remained unknown. Using biochemical approaches in yeast, we discovered that mechanochemical forces on the NC regulate CAT tailing. We propose CAT tailing initially operates in an “extrusion mode” that increases NC lysine accessibility for on-ribosome ubiquitination. Thr in CAT tails enhances NC extrusion by preventing formation of polyalanine, which can form α-helices that lower extrusion efficiency and disrupt termination of CAT tailing. After NC ubiquitylation, pulling forces on the NC switch CAT tailing to an Ala-only “release mode” which facilitates nascent chain release from large ribosomal subunits and NC degradation. Failure to switch from extrusion to release mode leads to accumulation of NCs on large ribosomal subunits and proteotoxic aggregation of Thr-rich CAT tails.

**One sentence summary:** Mechanochemical forces regulate the composition of CAT tails to extrude or release stalled nascent chains and recycle ribosomes.

## INTRODUCTION

Ribosomes can stall and collide during translation due to inefficient decoding of codons, structural impediments in mRNA, insufficient translation factors, or translation of poly(A) sequences past the stop codon. Stalled ribosomes block translation and produce incompletely synthesized nascent polypeptide chains (NCs) that can be nonfunctional or toxic to cells. The Ribosome-associated Quality Control (RQC) pathway dissociates and recycles stalled ribosomes and degrades stalled NCs (reviewed in references^1–4^). RQC first splits stalled ribosomes to form 60S-NC-tRNA ribonucleoprotein particles (RNPs) that are absent of 40S ribosomal subunits and mRNAs^5–9^ (Figure 1A). NCs then undergo two covalent modifications prior to release from the 60S ribosomal subunit: ubiquitylation by the E3 ligase Ltn1(yeast)/Listerin(human)^10,11^ and the addition of C-terminal amino acids in a conserved process called <u>CA</u>rboxyl-<u>T</u>erminal tailing, or “CAT tailing” initiated by the protein Rqc2(yeast)/NEMF(human)^12–16^. Rqc2p recognizes 60S-NC-tRNA RNPs and recruits specific charged tRNAs to the 60S A-site. Rqc2p then drives elongation of the NC with amino acids through non-canonical elongation independent of mRNA, the 40S ribosomal subunit, or energy input^12,17–21^. These C-terminal extensions are called “CAT tails”.

**Figure 1.**
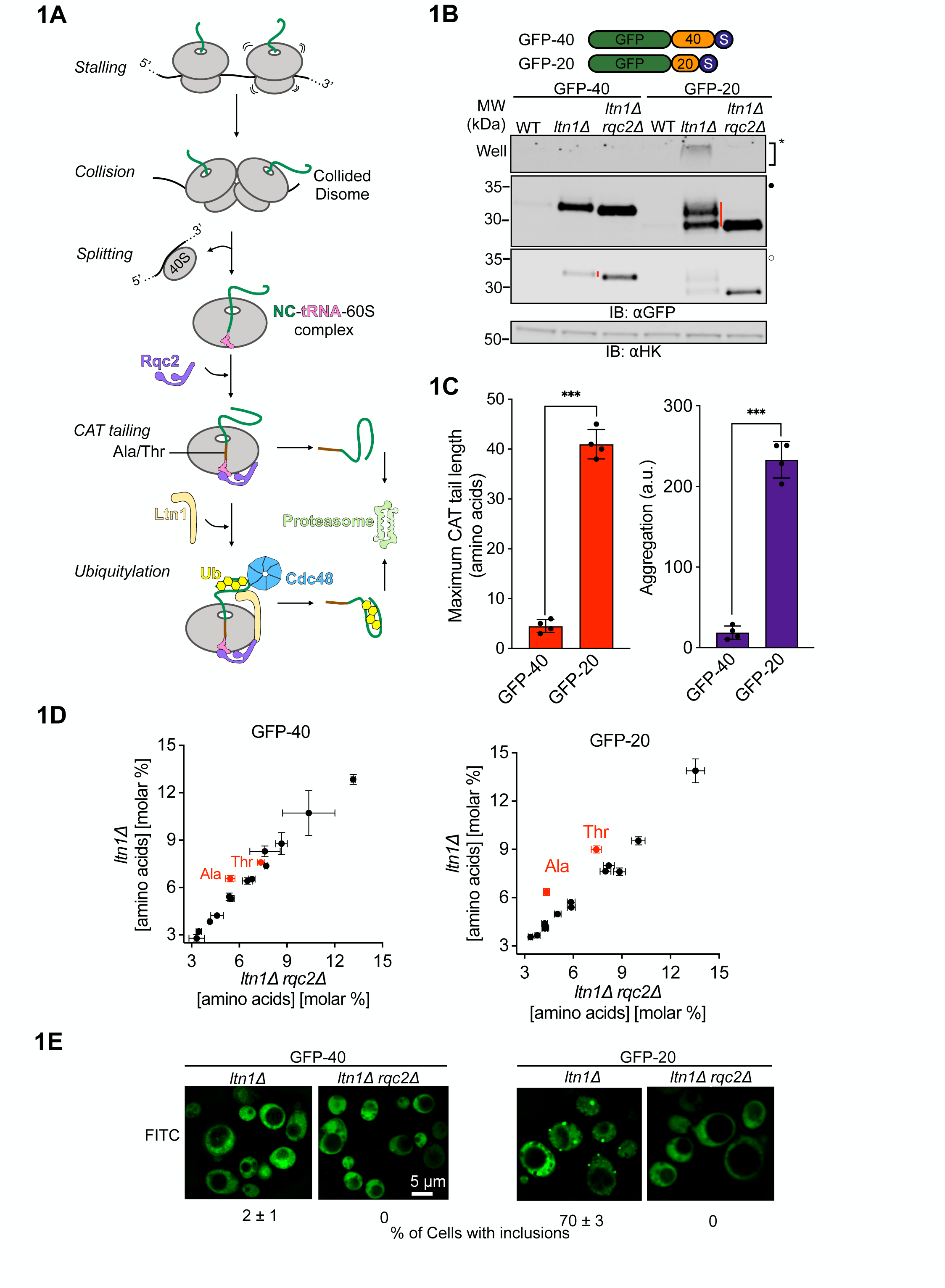
CAT tails have diverse composition, degradation, and aggregation propensities. **(A)** Schematic representation of the RQC pathway in yeast. Translational stalls cause ribosomes to collide and form disomes. Stalled disomes are recognized and the leading ribosome is split to generate a NC-tRNA-60S complex. Rqc2p recruits charged tRNA to the 60S as part of mRNA-free elongation of the NC with CAT tails. Ltn1p ubiquitylates the NC and Cdc48p extracts the ubiquitylated NC from the 60S ribosomal subunit. Both CAT tailing and ubiquitylation facilitate degradation of NC by the proteasome. Ub, ubiquitin. (**B)** Cartoon depicting two model RQC substrates, each fused to GFP via linkers (in orange) of lengths 20 and 40 amino acids followed by polyarginine stall sequence and stop codon (in navy blue). Whole cell immunoblots (IBs) of lysates containing model substrates expressed in different strains. SDS-insoluble aggregates near the well of the gel are indicated by asterisk (*). Filled (●) and empty (○) dots represent different contrast levels for the same image throughout the figures. Red vertical bars on the right side of bands indicate CAT tails. **(C)** Maximum CAT tail lengths were quantified by calculating the upshift in molecular mass between *ltn1Δ* and *ltn1Δrqc2Δ* strains and dividing that by the average amino acid mass, 110 Da. Protein aggregation was quantified by densitometric analysis of the areas surrounding the wells in the SDS-PAGE gel for each reporter in the *ltn1Δ* background, with the results normalized relative to the aggregation observed in the *ltn1Δrqc2Δ* strain. **(D)** Total amino acid analysis of substrates GFP-40 (*left)* and GFP-20 (*right)* from strains that are CAT tailing competent (*ltn1Δ*) and CAT tailing incompetent (*ltn1Δ rqc2Δ*). Error bars indicate s.e.m. from n=3 independent experiments. **(E)** Fluorescence microscopy of cells expressing model RQC substrates. Exposure durations for *ltn1Δ rqc2Δ* cells were longer than for *ltn1Δ* cells. Percentages of cells with visible GFP inclusions were calculated from n=3 independent experiments with ∼350 cells counted per experiment.

CAT tails are composed of alanine (Ala) in bacteria^13^, Ala and threonine (Thr) in yeast^12,22^, and Ala and other amino acids in *Drosophila* and humans^14,15,23^. In eukaryotes, CAT tails facilitate degradation of the NC by two parallel mechanisms. On the ribosome, CAT tails extrude the NC from the exit tunnel. This enhances the efficiency of Ltn1p in ubiquitylating NC by exposing any exit tunnel buried lysines^24^ and by increasing the accessibility of surface exposed lysines on structured NCs^22^. The NC must then be released from the 60S ribosomal subunit prior to degradation, as the topological restraint of the P-site tRNA and extra-ribosomal NC domain can anchor the NC to the ribosome^25^. The nuclease Vms1/ANKZF1 and peptidyl-tRNA hydrolases have been proposed to cleave the P-site tRNA and facilitate NC release^26–29^. Once off the ribosome, CAT tails act as C-terminal degrons that mark stalled NCs for proteasomal degradation, serving as a backup pathway in case Ltn1p does not successfully target the NC^22,30–32^ (Figure 1A). In bacteria, CAT tails function as degrons^13,18^ and have been proposed to enhance NC release from the large ribosomal subunit ^29^. In contrast to these beneficial cellular functions, CAT tailed proteins can also form cytotoxic protein aggregates that activate stress pathways and compromise cellular and organismal health^33–37^. Misregulation of CAT tailing has been proposed to explain neurodegeneration caused by mutations in NEMF in humans^38^ and phenotypes of Alzheimer’s and Parkinson’s disease models in flies^14,23,39,40^. Yet the relationship between CAT tail composition and function has remained unknown.

Here we discovered and studied diverse CAT tails on model substrates in yeast. Our findings suggest that cells regulate CAT tail composition to achieve two different functions. First, CAT tails comprising both Ala and Thr function in an “extrusion mode” to displace the NC out of the exit tunnel and increase accessibility of NC lysine residues for ubiquitination. Thr incorporation into CAT tails increases their efficiency of NC extrusion by preventing the formation of α-helices. Second, Ala-only CAT tails act in a “release mode” to facilitate release of the nascent chain from the 60S subunit and degradation by the proteasome. Switching from extrusion to release mode is driven by mechanochemical pulling forces on the NC which regulate CAT tail composition. These forces most prominently result from Cdc48p pulling on the ubiquitinated NC, but also folding of the NC outside of the exit tunnel, NC translocon-interactions, and electrostatic interactions between the NC and the exit tunnel interior. Threonine added during the extrusion mode of CAT tailing promotes efficient switching to the release mode when pulling force is applied to the NC. Failure to efficiently switch from extrusion to release mode leads to accumulation of NCs on ribosomes and proteotoxic aggregation of Thr-rich CAT tails.

## RESULTS

### CAT tails have diverse lengths, compositions, and aggregation propensities

In working with model substrates that constitutively induce ribosome stalling, we unexpectedly observed that different substrates had qualitatively different CAT tails (Figure 1B). CAT tails were visualized by comparing SDS-PAGE mobility in an *ltn1Δ* strain (to increase stalled NC detectability) to an *ltn1Δ rqc2Δ* strain (which cannot make CAT tails). A model substrate consisting of GFP followed by 20-amino acid linker and a stall-inducing polyarginine tract, GFP-20, was upshifted up to ∼3.5 kDa, corresponding to a CAT tail up to ∼35 amino acids in length (Figure 1B). By contrast, CAT tails on a similar substrate yet with a longer, 40 amino acid linker, GFP-40, were dramatically smaller (Figure 1B, 1C). To observe CAT tails in GFP-40 with more precision, we ran samples on a higher percentage acrylamide gel (Figure S1A). GFP-40 was uniformly upshifted by ∼0.5 kDa, even when Rqc2 was overexpressed, suggesting a CAT tail length of ∼4-5 amino acids that is unaffected by Rqc2p levels (Figure S1B). Similar observations were made in a WT (rather than *ltn1Δ*) background using model substrates that cannot be efficiently targeted by Ltn1p due to a lack of surface-exposed lysines (1K-sGFP-20 and 1K-sGFP-40)^41^ (Figure S1C-1D).

To determine the amino acid composition of CAT tails from GFP-20 and GFP-40, we immunopurified these proteins and performed total amino acid analysis. Consistent with our previous work^12^, CAT tails from GFP-20 contained roughly equal amounts of Ala and Thr (Figure 1D). In contrast, CAT tails from GFP-40 were composed of Ala while Thr enrichment was not observed. Analysis of the molar percentage of each amino acid suggested that CAT tails in GFP-40 were made up of ∼4-5 Ala. We conclude that GFP-20/1K-sGFP-20 and GFP-40/1K-sGFP-40 are CAT tailed in qualitatively distinct manners, with “long” Ala/Thr tails and “short” Ala CAT tails, respectively.

CAT tails have been found to aggregate and drive proteotoxic stress^12,33–35^. To evaluate the solubility of CAT tails produced in our model substrates, we compared levels of SDS-insoluble CAT tail protein aggregates (Figure 1B-1C) and localization by microscopy (Figure 1E). CAT tailed GFP-20/1K-sGFP-20 (Figure S1C) was less soluble and formed more cellular inclusions than GFP-40/1K-sGFP-40 (Figure S1D) in all backgrounds. Our results demonstrate that CAT tails on different stalled model substrates show a diversity of length, composition, and aggregation propensity, with GFP-40 appended with short, soluble CAT tails composed of Ala while GFP-20 received longer, Ala/Thr CAT tails that were aggregation prone.

### CAT tail composition determines degradation and aggregation

CAT tails function as “degrons” that, in parallel to ubiquitin added by Ltn1p, mark stalled NCs as substrates for degradation by the proteasome^22,30,31^. In an earlier study we showed that aggregation of CAT tails was anticorrelated with their ability to function as degrons^35^ and we here observed that levels of GFP-40 appeared lower than GFP-20 in an *ltn1Δ* background (Figure 1B). We therefore hypothesized that soluble short CAT tails would be more potent degrons than aggregation-prone long CAT tails. To quantitatively assess this, we generated internally controlled, ratiometric versions of our model substrates using RFP as an internal expression control followed by a tandem T2A peptide bond-skipping sequences and measured fluorescence by flow cytometry at steady state and after blocking synthesis with the translation inhibitor cycloheximide (CHX)^22^ (Figure 2A). Consistent with our hypothesis, the short CAT tailed substrate, GFP-40, accumulated in *ltn1Δ* cells at approximately half the level of the the long CAT tailed GFP-20 and was degraded at a rate 1.5-times faster than GFP-20 (Figure 2B).

**Figure 2.**
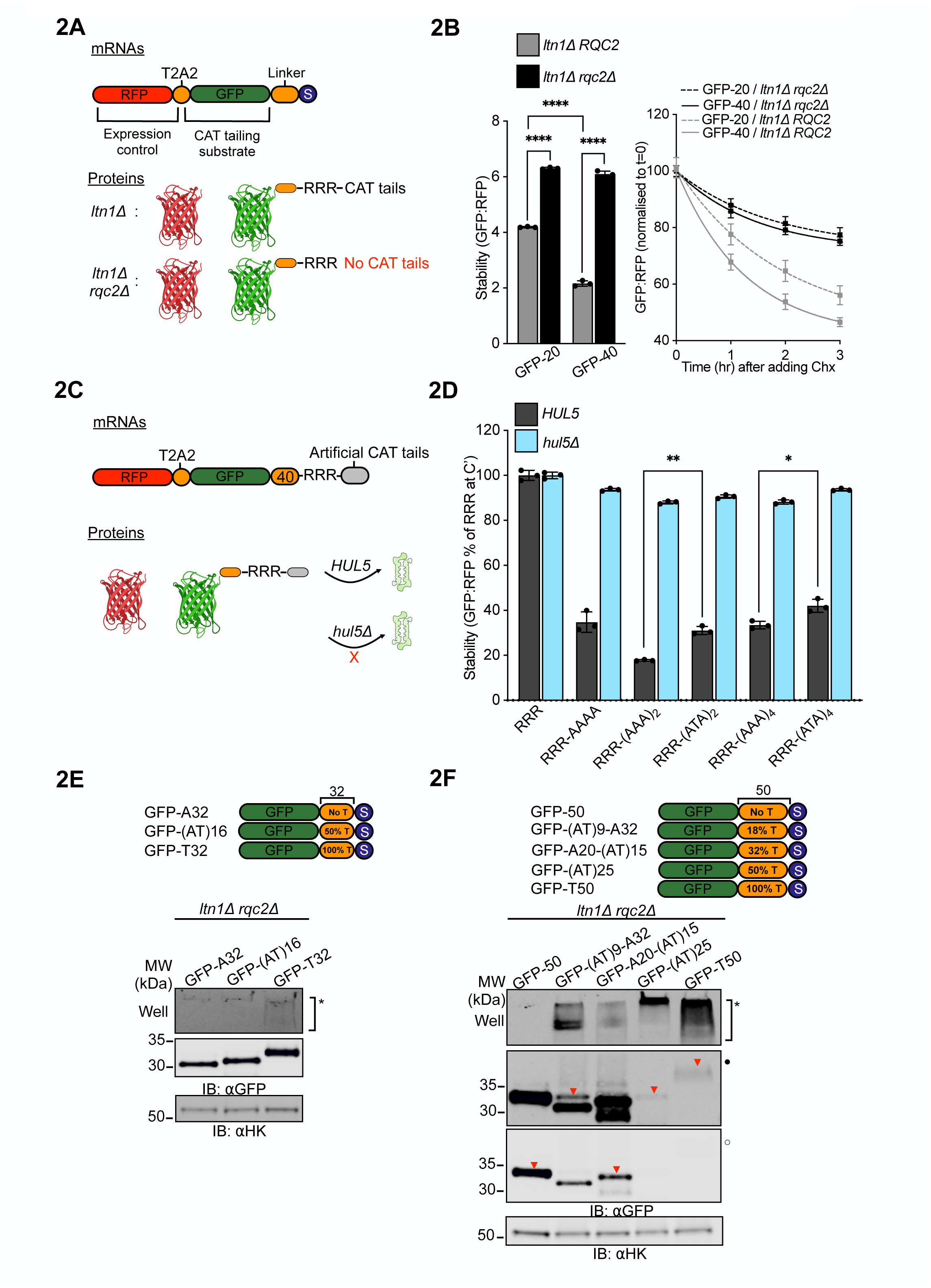
CAT tail sequence determines degradation and aggregation. **(A**) Schematic of expression controlled mRNA constructs and expected protein products used in assays for steady state measurement of stability and cycloheximide chases. **(B)** *Left*, stability measurements for GFP-20 and GFP-40 substrates in cells with intact *RQC2* (*ltn1Δ*) and lacking *RQC2* (*ltn1Δ rqc2Δ*); data are reported as mean values. *Right*, normalized stability measurements for GFP-20 and GFP-40 in the *ltn1Δ* and *ltn1Δ rqc2Δ* backgrounds, over a 3 hour chase following a 200 μg/mL cycloheximide pulse. \*\*\*\**P* < 0.0001, *P* values are derived from a two-tailed *t*-test. **(C)** Schematic of expression controlled mRNA constructs and expected protein products with hard coded CAT tails used in assays measuring stability in WT and *HUL5*-deficient (*hul5Δ*) cells. **(D)** Stability measurements of hard coded CAT tail constructs with the indicated C-terminal sequences in WT and *hul5Δ* cells. Data are presented as mean values, normalized to the construct containing only the RRR sequence in WT cells. Error bars indicate s.e.m. from at least three independent experiments (n ≥ 3). Stability levels between pairs of similar sized hard coded CAT tails in WT cells were compared via Student’s *t* test. \*\**P* < 0.01; \**P* < 0.05. R, arginine; A, alanine; T, threonine. **(E-F)** Cartoon depicting substrates with two distinct linker lengths, 32 and 50, each containing varying percentages of threonine residues (indicated as a proportion of total amino acids in the linker). Whole cell IBs of lysates prepared from non-CAT tailing *ltn1Δ rqc2Δ* cells expressing the indicated substrates. Inverted red triangles (▾) indicate the expected position for protein bands in SDS-PAGE/IBs.

To test the effects of reduced CAT tailing on GFP-20 and GFP-40 phenotypes without completely blocking CAT tailing, we leveraged an Rqc2 mutant, *rqc2*-D9A, that exhibits a partial loss of function phenotype in CAT tail synthesis^22^. Consistent with this, we observed a reduction in CAT tail length and complete loss of aggregation in the *rqc2*-D9A background (Figure S2A, 2B). No CAT tailing or aggregation was observed in the Rqc2-null strain, as previously observed^12,22^, or in the catalytically dead *rqc2*-D98A mutant. Analysis of ratiometric reporters in *rqc2*-D9A revealed a modest increase in steady state levels (significant for GFP-40, not significant for GFP-20) (Figure S2C) and decrease in degradation rate via CHX chase assay (Figure S2D) for both GFP-40 and GFP-20 reporters, suggesting that degron function was moderately attenuated in *rqc2*-D9A. Thus, partial impairment of CAT tailing by *rqc2*-D9A leads to decreased CAT tail length, decreased degron function, and abolished aggregation.

In mammalian cells, C-terminal Ala tails function better as degrons than Ala/Thr tails^31,42^. We also found this to be the case in yeast, with six C-terminal Ala being optimal among the sequences we tested for targeting substrates to the proteasome via the ubiquitin ligase Hul5p (previously shown to target CAT tails off the ribosome^22^). Inclusion of Thr lowered degron efficiency (Figure 2C-D). To investigate if CAT tail sequence explains the difference in aggregation propensity we observed in GFP-20 and GFP-40, we fused GFP with 32-amino acid “hard coded” CAT tails containing only Ala, only Thr, or randomly mixed Ala and Thr. Hard coded CAT tails that contained only Thr exhibited partial aggregation, whereas no aggregation was observed for Ala-only or mix of Ala/Thr extensions at this length. (Figure 2E). We then fused GFP with longer 50-amino acid hard coded CAT tails containing differing proportions of Ala and Thr and found aggregation increased with Thr content. This is consistent with studies where inclusion of Thr in prion and α-synuclein proteins increased aggregation propensity^43–45^. Raising Thr content to 50% or more resulted in nearly complete protein aggregation (Figure 2F). Together, these results suggest that both length and Thr content of CAT tail sequences explain the differences in degradation and aggregation properties in GFP-20 and GFP-40, with short Ala tails functioning as effective, soluble degrons and longer Thr containing CAT tails acting as poor degrons with increased aggregation propensity.

### Mechanical forces regulate CAT tail composition

We next investigated the mechanism explaining differences in CAT tailing we observed between substrates. Vms1p is a protein that has been shown to antagonize CAT tailing by competing with Rqc2p binding^27,46,47^ and cleaving the P-site tRNA to release ubiquitylated NC^28,48^. Deletion of the *VMS1* gene had no effect on CAT tail lengths of our substrates in wild type and *ltn1Δ* backgrounds, although aggregation propensity was reduced for some substrates (Figure S3A). We therefore considered what intrinsic features of our model substrates may contribute to different CAT tail phenotypes. One basic feature of the stalled NC that is predicted to be different between GFP-20 and GFP-40 is the folding state. In GFP-40, the GFP sequence fully emerges from the ∼40 amino acid long ribosomal exit tunnel and can fold into its native state^49^. By contrast, the GFP-20 substrate has 20 amino acids sequestered within the exit tunnel, which would be predicted to prevent its folding. NC folding has been demonstrated to generate a pulling force that extends all the way to the P-site tRNA^50–53^ and regulates canonical translation^51,54–57^. We hypothesized that pulling force on the NC regulates CAT tail composition and termination.

To explore this hypothesis, we designed RQC substrates with different linker lengths. Our prediction was that CAT tailing would terminate when the full length GFP sequence emerged outside the ribosome and could thus fold and create a pulling force. That is, the length of CAT tail and the linker length were predicted to be inversely related. Consistent with this hypothesis, maximum CAT tail length plus the linker was estimated to be between 50-60 amino acids for all constructs, lengths at which the full GFP sequence would emerge from the ribosome and fold (Figure 3A-B). Aggregation generally correlated with CAT tailing (Figure 3C). Yet when linker length was increased to 50 amino acids, a length at which pulling is predicted to be reduced because the folded NC no longer abuts the ribosome^51,58^, aggregation of CAT tailed protein resumed (Figure 3A-C). A similar pattern of CAT tailing and aggregation was observed in analogous experiments where GFP was replaced by maltose binding protein (MBP)^59^, suggesting the effects observed were related to any NC folding rather than specific to GFP. Like GFP-50, MBP-50 showed increased aggregation over the 40-amino acid linker counterparts. MBP-70 had longer CAT tails and more aggregation than MBP-40 and MBP-50 (Figure S3B-D).

**Figure 3.**
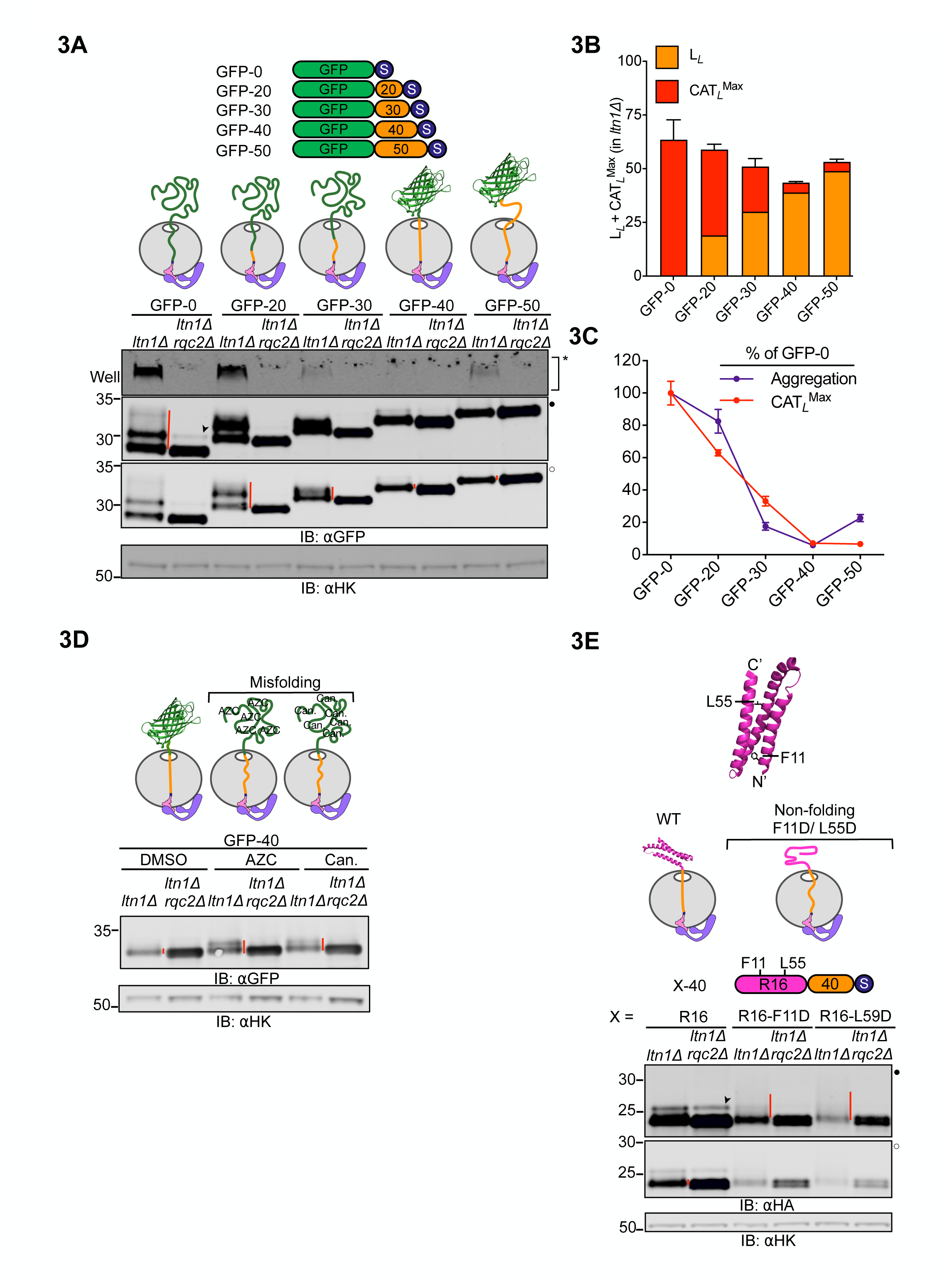
Folding-induced mechanical forces on the NC determine CAT tail composition. **(A)** Cartoon of RQC substrates with linkers of varying lengths (0, 20, 30, 40 and 50 amino acids) between GFP and a stalling sequence. The diagrams above IBs illustrate the predicted RNP-Rqc2p complex folding state for each RQC substrate in the stalled, pre-CAT tailing state. Arrowheads (➤) indicate readthrough products formed by bypassing the polyarginine arrest sequence. **(B)** Stacked bar graph showing the sum of linker and estimated maximum CAT tail length for each substrate in the *ltn1Δ* background. **(C)** Plot depicting the relative aggregation and maximum CAT tail length for each RQC substrate, normalized to the GFP-0 substrate, in the *ltn1Δ* background. Error bars indicate s.e.m from three biologically independent experiments. **(D)** *Top*, cartoons depicting the folded states of stalled NCs as part of the RNP-Rqc2p complex. Cells expressing GFP-40 were grown in media supplemented with DMSO (control), translation inhibitors, 1 mM AZC, or 200 μM canavanine for 3 hours to induce misfolding. *Bottom*, IBs showing changes in CAT tail lengths. **(E)** IBs of cells expressing RQC substrates containing WT and mutant versions of spectrin R16 protein. *Top,* structure of *E. coli* protein spectrin R16 showing the location of non-folding mutations (PDB: 5M6S). *Middle*, cartoon depicting RNP-Rqc2p complex with WT and misfolded spectrin R16 in the stalled, pre-CAT tailing state. *Bottom*, IBs showing CAT tails observed for WT, F11D and L55D variants of spectrin R16 with a 40-amino acid linker and R12 stalling sequence.

We next tested the pulling-force model for CAT tail regulation with four additional, orthogonal approaches. First, we sought to chemically modulate cotranslational protein folding by growing cells in media supplemented with unnatural amino acid analogs that disrupt protein structure. Consistent with this hypothesis, GFP-40 showed increased CAT tail lengths, up to 20-25 amino acids, in the presence of *L*-Azetidine-2-carboxylate (AZC), a proline analog, or canavanine, an arginine analog (Figure 3D). Corresponding experiments performed with the 1K-sGFP-40 reporter in *LTN1*-intact cells yielded similar results whereas no increase in CAT tail lengths were seen for the GFP-20/1K-sGFP-20 reporters (Figure S3E).

In a second approach to modulate NC protein folding we used spectrin R16, a protein for which cotranslational folding has been characterized^60,61^, and introduced independent point mutations that produce defects in its cotranslational folding. All constructs featured an identical 40-amino acid linker to comparably ensure they cleared the ribosome before stalling. As predicted, wild type, folding-competent spectrin R16 formed short, 0.5 kDa (∼5 amino acid) CAT tails in *ltn1Δ* whereas both non-folding mutant versions produced smears indicative of longer CAT tails of up to 6.5kDa (∼65-70 amino acids) (Figure 3E).

We next tested if the effect of forces generated during cotranslational insertion of the NC into the endoplasmic reticulum (ER) lumen^62–65^ caused early termination of CAT tailing (Figure 4A). We appended a carboxypeptidase Y-derived ER signal peptide to the N termini of GFP-20 and GFP-40 to generate spGFP-20 and spGFP-40. As predicted, spGFP-20 had ∼2.5 times greater population of short CAT tails than GFP-20 in the *ltn1Δ* background (Figure 4B-C) whereas spGFP-40 remained unaffected (Figure S4A). These phenotypes remained qualitatively the same after deletion of *VMS1* (Figure S4B), suggesting that CAT tail termination was not driven by Vms1p at the ER.

**Figure 4.**
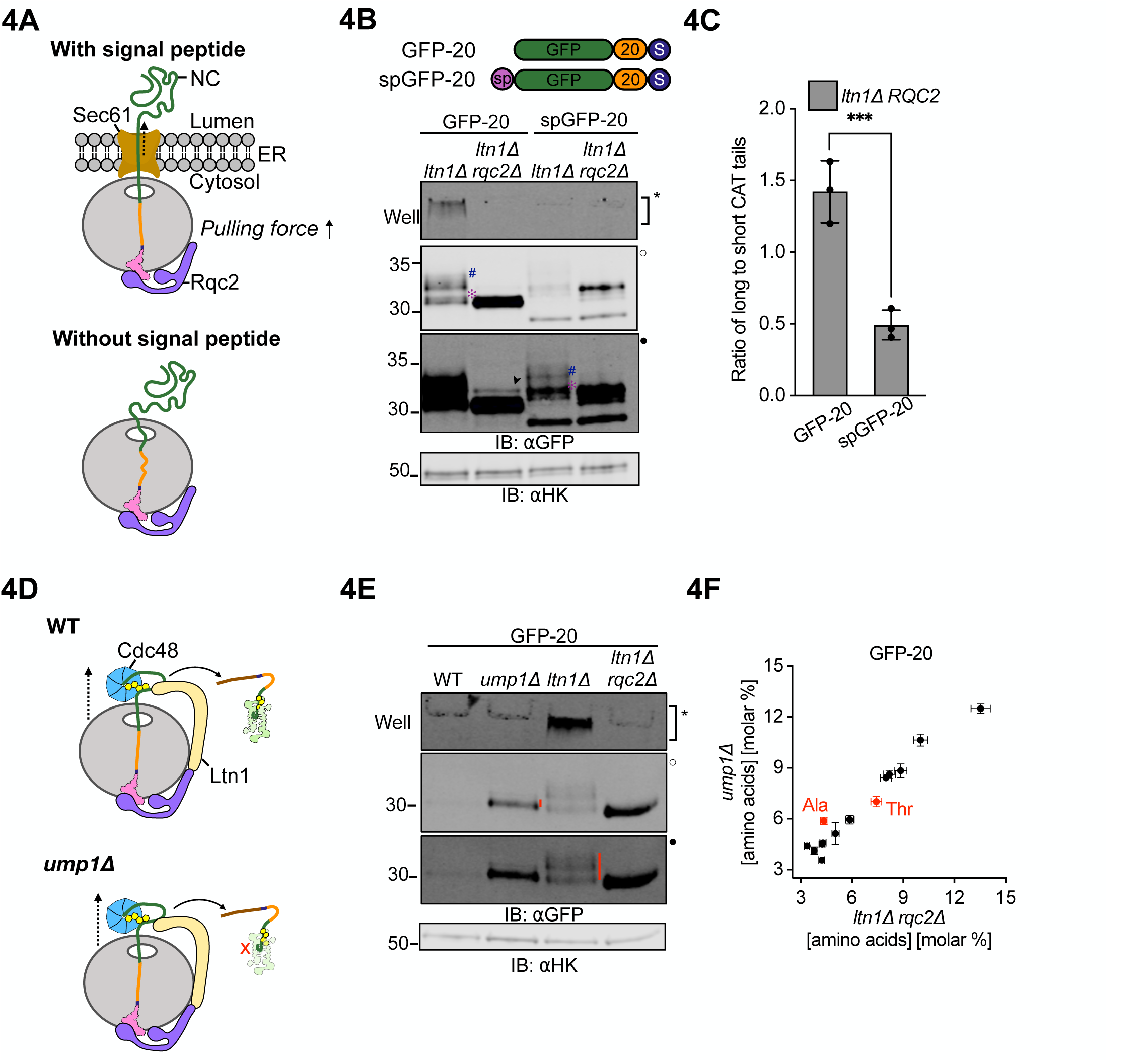
Extrinsic mechanical forces on the NC determine CAT tail sequence. **(A)** Schematic showing the organization of a NC-tRNA-60S-Rqc2p at the Sec61 translocon complex at ER (*top*) and in the cytoplasm (*bottom*). Dotted arrow indicates the direction of the pulling force. **(B)** Cartoons of ER-directed RQC substrate spGFP-20 with an N terminal signal peptide derived from carboxypeptidase Y, alongside cytosolic RQC substrates (no signal peptide). Cells expressing these RQC substrates were analyzed by immunoblot. Hash (#) and asterisk (✻) signs next to CAT tails denote long and short CAT tails respectively. Arrowhead (➤) indicates readthrough product formed after bypassing the polyarginine arrest sequence. **(C)** Densitometric analysis was used to quantify long and short CAT tailed species and are represented as a ratio (i.e. # to ✻) (See Supplementary File S2). Error bars indicate s.e.m. from n=3 independent experiments.\*\*\**P* <0.001; Student’s *t*-test. **(D)** Schematics showing the RQC role of *CDC48* in different strain backgrounds. *Top*, in WT cells Cdc48p binds and extracts the ubiquitylated NC from the 60S for proteasomal degradation. *Bottom*, cells without the *UMP1* gene exhibit impaired proteasome assembly and this stabilizes NCs that have been ubiquitylated by Ltn1p and pulled by Cdc48p. **(E)** GFP-20 reporter expressed in *ump1Δ* cells alongside WT, non-ubiquitylated (*ltn1Δ*) and non-CAT tailed (*ltn1Δ rqc2Δ*) controls. **(F)** Total amino acid analysis of immunoprecipitated GFP from *ump1Δ* cells and *ltn1Δ rqc2Δ* cells expressing GFP-20. Error bars indicate s.e.m. from n=3 independent experiments.

Previous work identified the Cdc48 ATPase complex as an RQC factor and proposed that it physically extracts ubiquitylated stalled NCs^5,7,66^. In a final approach to alter forces pulling on the NC, we compared CAT tailing in strains with reduced Cdc48p activity, predicting longer CAT tails when pulling by Cdc48 was reduced. We first measured CAT tails of the GFP-20 and GFP-40 substrates in W303 cells with wild type *CDC48* and a temperature sensitive mutant, *cdc48-3*, with constitutive RQC defects at the semi-permissive temperature of 30°C^66^ (Figure S4C-D). This experiment was performed in the W303 strain background (to access the *cdc48-3* mutant) unlike other experiments in the study, which were performed in the BY4741 strain background. Cells were treated with bortezomib to block degradation of ubiquitylated CAT tailed proteins and enhance their detection. As predicted, a banding pattern for *cdc48-3* that was consistent with longer average CAT tails was observed for GFP-20 but not GFP-40, which folds and therefore can generate a NC pulling force without Cdc48p. Moreover, deleting *LTN1* in *cdc48-3* cells resulted in more aggregates but only a minor change in banding pattern for GFP-20, suggesting that the effect of *LTN1* deletion on CAT tails occurred through loss of Cdc48p function. However, the putative CAT tail band in GFP-20 that increased in relative intensity in *cdc48-3* migrated at or near the same size as the readthrough product observed in the *rqc2Δ* background, complicating interpretation of the results. We then took an orthogonal approach to block Cdc48p activity by comparing CAT tails in a WT strain to those without *LTN1*, which is required for NC ubiquitylation and thus recruitment of Cdc48p^5,7^. To increase observability of the model substrates when *LTN1* was intact, we crippled the proteasome by deleting the proteasome maturation factor Ump1p^67^(Figure 4D). Consistent with our model, CAT tails of GFP-20 (Figure 4E) but not GFP-40 (Figure S4E) were dramatically shorter in *ump1Δ* (Cdc48p recruited to the NC) compared to *ltn1Δ* (Cdc48p not recruited to the NC). Amino acid analysis of GFP-20 revealed that the CAT tails in *ump1Δ* were composed only of Ala (Figure 4F), while the CAT tails observed in the *ltn1Δ* background were composed of both Ala and Thr (Figure 1D, right panel). GFP-20 in *ump1Δ* had the same CAT tail composition found on GFP-40 in an *ltn1Δ* background (Figure 1D, left panel), which folds and therefore, like GFP-20 in *ump1Δ,* experiences NC pulling forces. Together, our results support a model where tension generated by pulling forces on stalled NCs (via Cdc48p, NC folding, or membrane translocation) modulates CAT tail composition and length. When no pulling is present, both Ala and Thr CAT tails are added. Pulling drives addition of Ala-only CAT tails and termination of CAT tailing.

### NC-60S ribosome interactions regulate CAT tail composition

While our data is consistent with a hypothesis that pulling force on the NC regulates CAT tailing, it is also consistent with a model where the extra-ribosomal NC folding state is sensed (perhaps by a chaperone) and regulates CAT tailing. To distinguish between these models, we sought to create a NC pulling force within the exit tunnel that does not affect the extra-ribosomal NC. Mechanochemical forces on the NC occur if the NC has electrostatic interactions with the negatively charged ribosomal exit tunnel^68–70^. In our experiments so far we used linkers composed of a mix of neutral and charged amino acids. To test how NC/exit tunnel interactions affect CAT tailing, we introduced charged tracts in GFP ∼15 amino acids away from the PTC in a “constriction site” where electrostatic interactions within the ribosome exit tunnel are most pronounced^70–72^ (Figure 5A). Strikingly, encoding positive charges via five consecutive lysines sharply reduced CAT tailing and abolished aggregation in GFP-0. When a negatively charged aspartate tract was introduced to GFP-0, CAT tailing and aggregation remained (Figure 5B). In a symmetric experiment, an aspartate tract caused a sharp increase in Rqc2-dependent aggregation of GFP-40 (Figure 5C). We then supplanted seven amino acids near the constriction site of a previously characterized linker that is stretched end to end in the ribosome when stalled^54,73^. Substituting with negatively charged residues led to dramatic increases in CAT tail length and aggregation (Figure 5D). Taken together, our data suggest that mechanochemical forces on the NC regulate CAT tailing. NC pulling from outside the ribosome match the effect of NC-exit tunnel attraction and produce short Ala CAT tails, while no NC pulling matches the effect of NC-exit tunnel repulsion and increases the length, aggregation, and Thr content of CAT tails.

**Figure 5.**
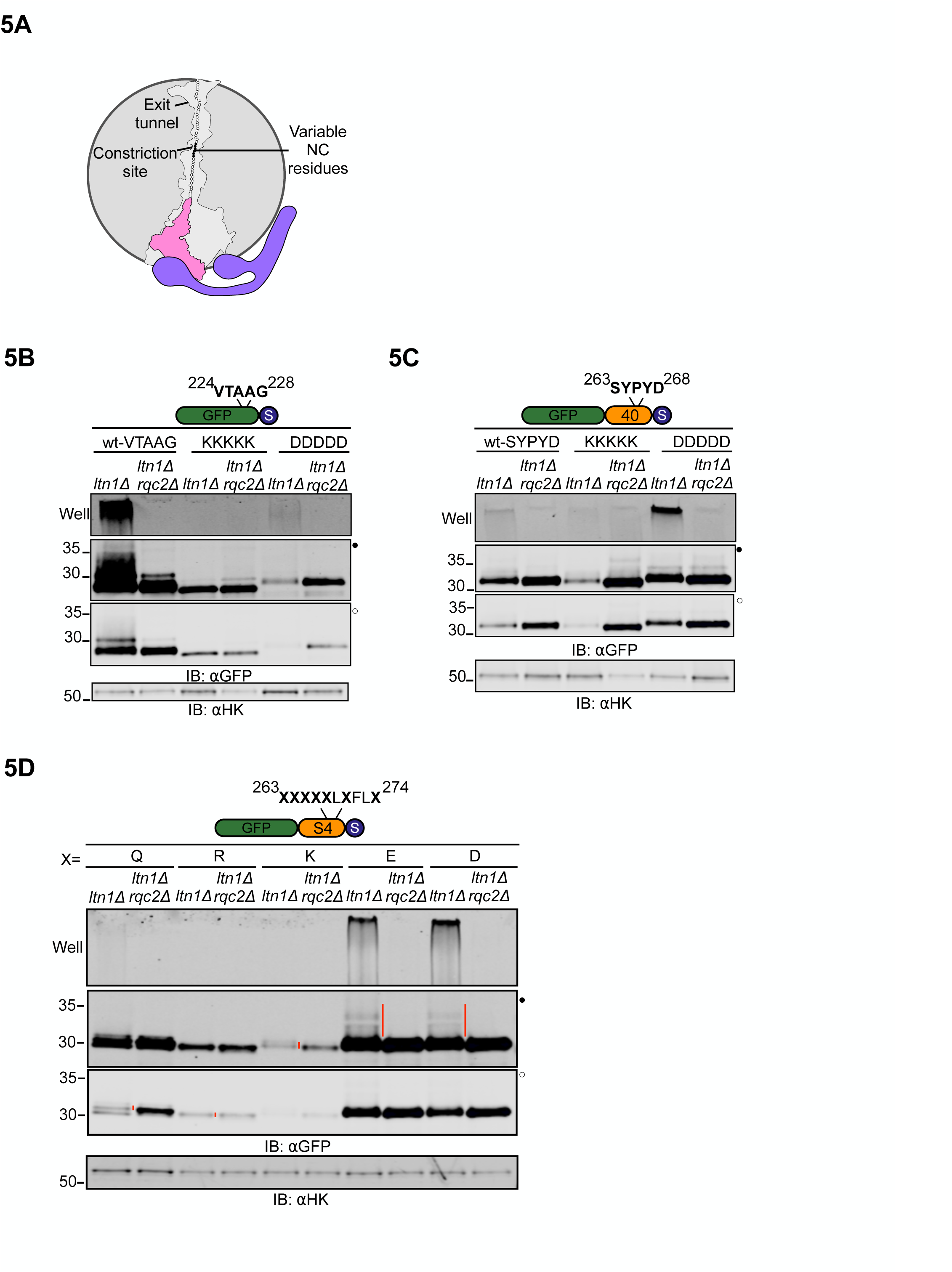
NC-60S exit tunnel interactions regulate CAT tail sequence. **(A)** Schematic of the 60S ribosome exit tunnel showing NC residues chosen for mutagenesis (in black) and the constriction site; outline of exit tunnel based on PDB: 5GAK^90^. **(B)** Residue 224 of GFP-0 reporter, situated 15 amino acids upstream of stalling sequence, to residue 228 were mutated to five consecutive Lys or Asp and analyzed by immunoblots. **(C)** Residue 263 to 268 of GFP-40 reporter were mutated to Lys or Asp as indicated and analyzed by immunoblots. **(D)** Immunoblots of RQC reporter consisting of GFP and the polyarginine stalling sequence linked by a 40 amino acid sequence (S4) derived from the N terminus of the fourth transmembrane segment of a bacterial voltage-gated potassium channel. This peptide sequence is reported to be fully extended inside the ribosome exit tunnel 54^54,73^. Residue 263 was situated 15 amino acids upstream of the stalling sequence and residues indicated by X were mutated to Gln, Arg, Lys, Glu or Asp, as indicated.

### Thr increases the efficiency of NC extrusion by CAT tails

A proposed function of CAT tails is to enhance Ltn1p’s ability to ubiquitylate lysines on NCs by extruding the NC from the exit tunnel and thus increasing the accessibility of exit tunnel-occluded lysines^24^ and lysines on structured regions of NCs^22^. Since our amino-acid analysis data suggested that Thr is only present in CAT tails prior to NC ubiquitylation and pulling by Cdc48p, we hypothesized that Thr plays a functional role in extrusion. Among all amino acid homopolymers, polyalanine exhibits the highest propensity for forming ⍺-helices within the ribosome exit tunnel^74–76^, which compacts the NC by up to ∼2x^73^. We hypothesized that ⍺-helix formation would compact CAT tails, rendering Ala-rich CAT tails poorly suited to extrude the NC from the exit tunnel and that Thr incorporation disrupts these ⍺-helices. Thr disruption of polyalanine ⍺-helices was supported by theoretical predictions of protein secondary structure (Figure 6A). To experimentally test this, we encoded hard coded Ala linkers of lengths 6, 13, 20, and 32 amino acids in a GFP-based stalling reporter that were predicted to form ⍺-helices. Since the exit tunnel is ∼40 amino acids long, these RQC substrates are unfolded when they stall and should be CAT tailed until the GFP is fully extruded and folds, generating pulling forces that terminate CAT tailing. Remarkably, despite the progressive increase in the number of linker residues, the CAT tails on all Ala-linker substrates were up to 5 kDa (∼50 amino acids)(Figure 6B-C), which is approximately the same maximum length we observed on GFP-0. By contrast, both Thr32 and (AlaThr)16 linkers resulted in CAT tails with maximum length 2 kDa (∼20 amino acids) (Figure 6C), consistent with a total CAT plus linker length of ∼40-50 amino acids as observed for GFP-based substrates without helix-forming linkers (Figure 3A and S3B). The increase in CAT tail length when a polyalanine linker was used was beyond what is explained by a 2x compaction of polyalanine due to its ⍺-helical structure. The presence of long polyalanine tracts in the ribosome exit tunnel may thus cause a failure to terminate CAT tailing upon folding of the extruded GFP. To test if long polyalanine tracts also cause failure to terminate CAT tailing under the pulling force of Cdc48, we expressed our reporters in cells in which Cdc48 pulling was intact but proteasome activity was reduced (via deletion of UMP1) in order to allow the observation of CAT tails (Figure 6D). While 13 or 20-Ala linkers gave the expected short CAT tail lengths that should result from NC pulling by Cdc48 (Figure 6D), a 32 Ala linker yielded a long CAT tail, an effect that was abolished when the linker included Thr (Figure 6E). These results suggest that extensive polyalanine in the ribosome exit tunnel prevents the termination of CAT tailing upon nascent chain ubiquitylation and pulling by Cdc48. Taken together, these observations are consistent with Thr in CAT tails increasing the efficiency of extrusion and allowing CAT tail termination when pulling forces are applied to the NC, an effect that likely happens because Thr disrupts the formation of polyalanine ⍺-helices. We term this form of CAT tailing “extrusion mode”.

**Figure 6.**
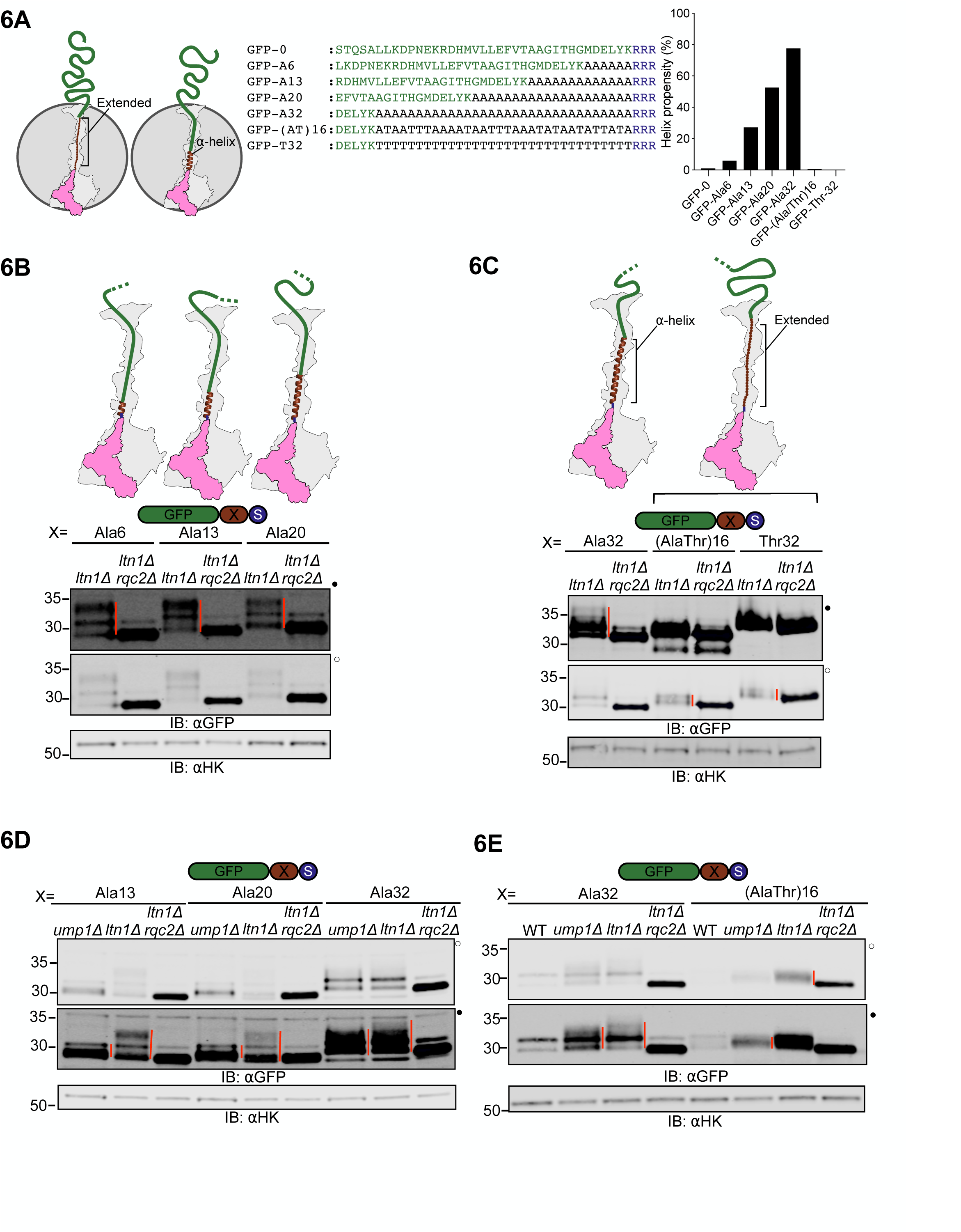
Thr in CAT tails increases the efficiency of NC extrusion and termination of CAT tailing upon NC pulling. **(A)** *Left*, schematic representation of the 60S ribosomal subunit showing the ribosome occluded portion of the NC in extended and ⍺-helix forms. In the extended form, the NC is stretched end-to-end. Formation of an ⍺-helix compacts the NC inside the ribosomal exit tunnel, retracting the extra-ribosomal portion of the NC into the ribosome. *Center*, sequences of the constructs used in experiments below depicting forty C terminal amino acids of RQC reporters before CAT tailing. *Right*, the helix forming propensity of the forty C terminal amino acids are depicted, as calculated from AGADIR algorithm^91^ using these parameters: temperature, 298K; ionic strength, 0.2M; and pH 7.0. **(B-C)** Reporters expressed in *ltn1Δ* and *ltn1Δrqc2Δ* cells and analyzed by immunoblots, with schematics above each reporter showing the predicted structure of hard coded linkers (extended or ⍺-helix) prior to CAT tailing **(D-E)** Reporters expressed in an *ump1Δ* background in addition to *ltn1Δ* and *ltn1Δ rqc2Δ*, analyzed by immunoblot as above.

### CAT tailing regulates NC retention and release from ribosomes

Our data suggested that Ala-only CAT tails are added when the NC is pulled. This represents the final stages of RQC, after the NC has been ubiquitylated and should now be released and degraded. Ala CAT tails have been proposed to aid NC release from ribosomes in bacteria^29^. We hypothesized that the short, Ala CAT tails observed on GFP-40 facilitate NC release in yeast.

To test our hypothesis, we quantified NC retention on different ribosomal subunits though fluorescent polysome profiling which can detect both RNA and fluorescent GFP levels, allowing normalized levels (by virtue of rRNA detection) of ribosome-bound fluorescent model substrate NCs to be quantified^77^. The polysomes were run in low salt conditions under which free 40S and 60S ribosomes associate into 80S ribosomes in vitro unless association-blocking factors like Ltn1, Rqc2, and eIF6 are present^11,78^. This allowed us to assess if NC were free from RQC factors (in the reassociated 80S fraction) or likely bound by RQC factors (in the 60S fraction). We predicted that release of GFP-40 from ribosomes would be aided by CAT tails. Consistent with this, we observed ∼3.5 fold increase of 60S- and 80S-associated GFP-40 fluorescence when CAT tailing was impaired (Figure 7A-B). No 60S or 80S fluorescence peaks were observed in a non-fluorescent background or strains expressing GFP-stop, demonstrating that stalling was required to accumulate NCs on ribosomes (Figure S7A). Performing the polysome gradient in high salt conditions that do not allow 40S/60S reassociation moved the GFP-40 peak almost entirely to the 60S fraction, showing that the 80S peak of GFP-40 in low salt was the result of *in vitro* recombination of 60S-GFP-40 complexes that do not include RQC factors that block reassociation (Figure S7B). Next, we tested if Thr-rich, aggregation prone CAT tails have a release function. We utilized the GFP-S4(D) substrate, which has a 40 amino acid aspartate-rich linker that yields long CAT tails that promote aggregation (Figure 5D). This substrate was not quantifiable on 60S ribosomes due to a large adjacent peak, likely originating from high molecular weight aggregates, however the 80S peak, which is a product of *in vitro* recombined 60S-NC complexes, could be quantified. GFP-S4(D) was at least ∼3 fold enriched on 80S in the WT background compared to GFP-40 and GFP-S4(K) (both of which garner short CAT tails) (Figure 7C). Blocking CAT tail synthesis had no effect on association of GFP-S4(D) with ribosomes. These data suggest that Thr-rich CAT tails do not promote NC release from 60S ribosomal subunit. We conclude that Ala but not Thr CAT tails promote release of the NC from the 60S-NC complex.

**Figure 7.**
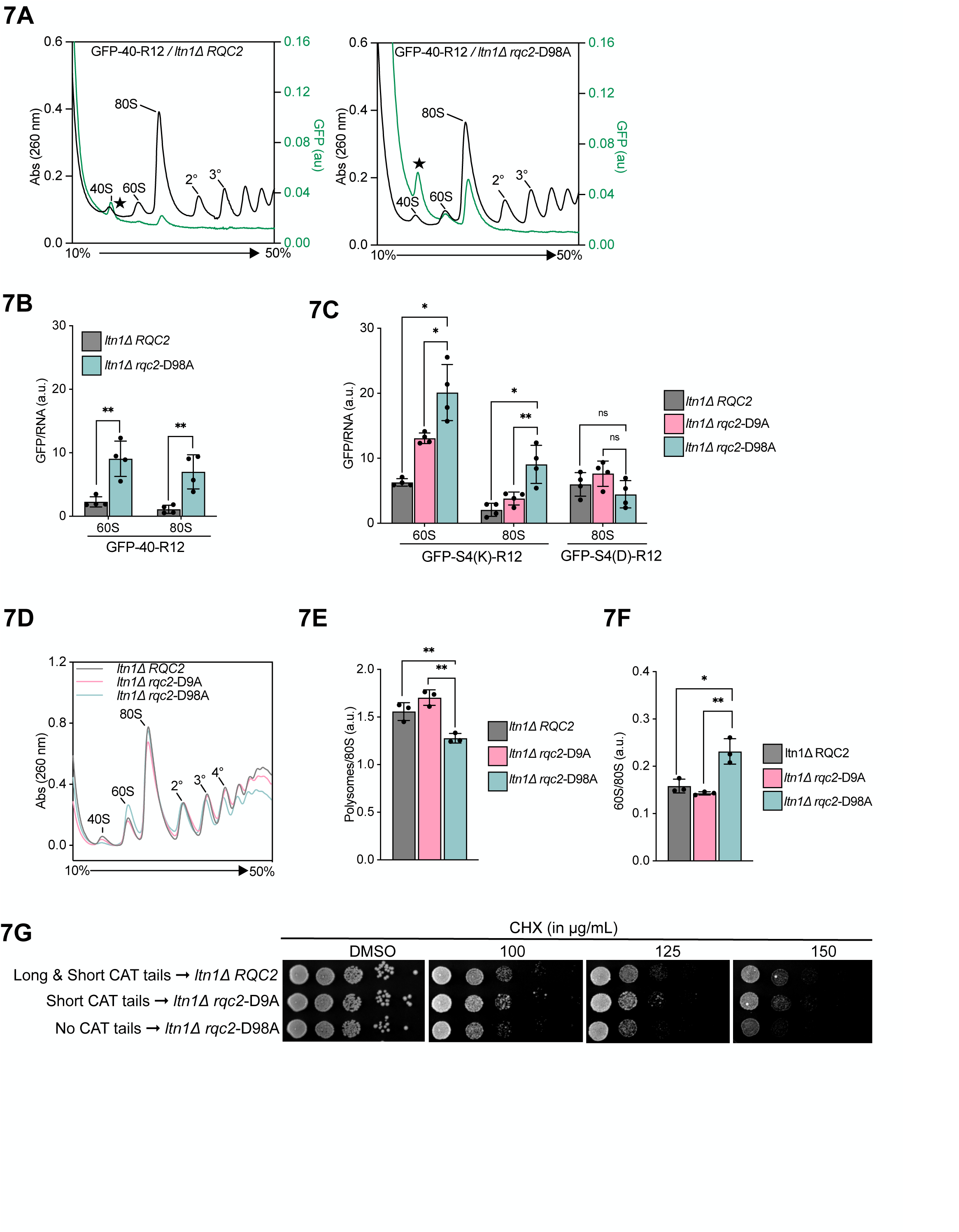
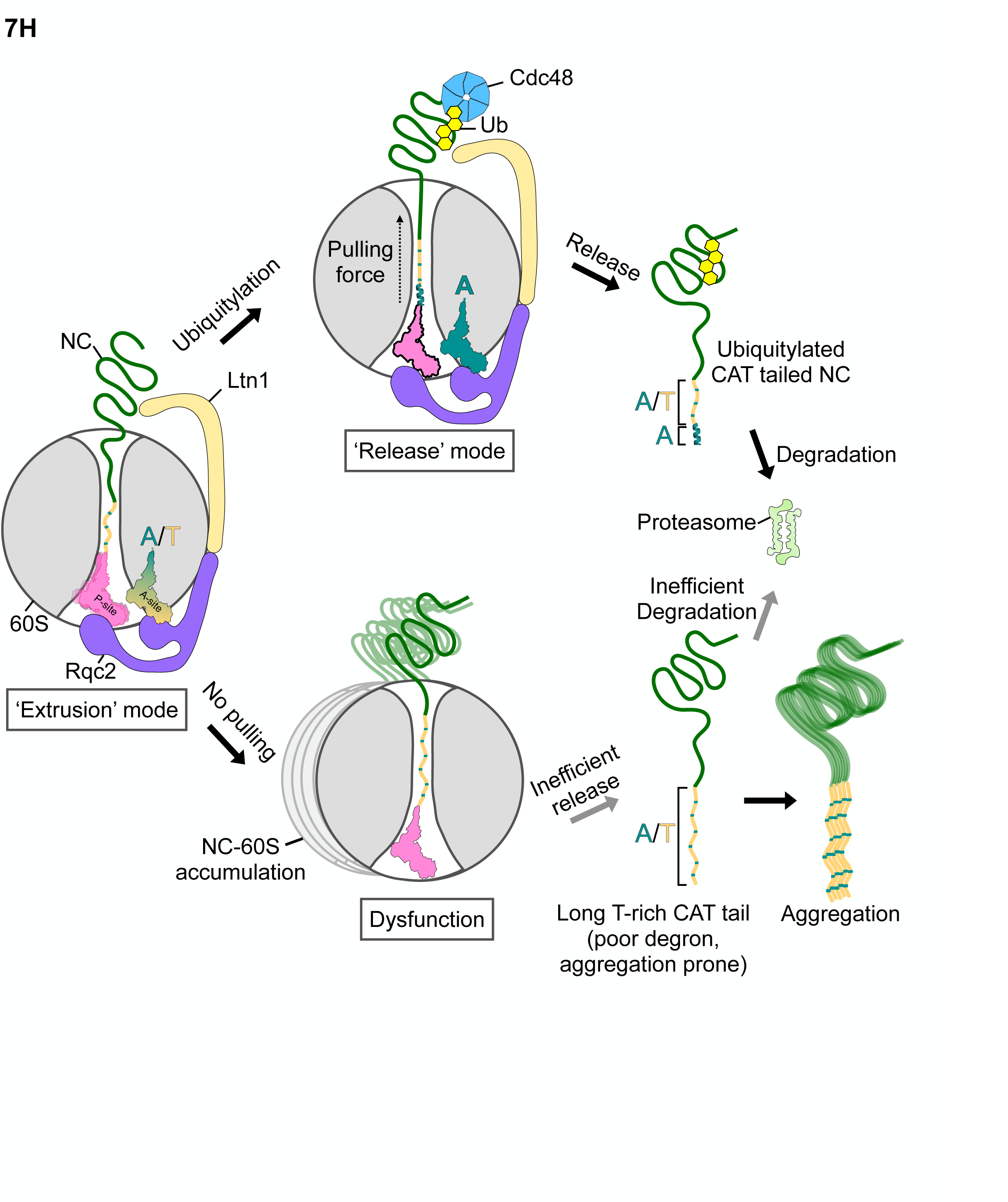
CAT tails regulate NC retention and release. **(A)** Sucrose gradients of the GFP-40 reporter expressed in *ltn1Δ* cells with WT *RQC2* and *rqc2*-D98A were analyzed by fluorescent polysome profiling. A background fluorescence peak at the 40S position is marked with a star (★) where visible (See Supplementary Figure S7A). **(B)** From the GFP fluorescence and RNA absorbance plot for each reporter/strain pair in **(A)**, the areas covered by 60S and 80S local peaks were calculated (See methods). Error bars in computed values indicate s.e.m. from four independent experiments. \*\**P* < 0.01, Student’s *t* test. **(C)** Similar quantification was performed for GFP-S4(K) and GFP-S4(D) reporters expressed in *ltn1Δ* cells with WT *RQC2*, *rqc2*-D9A and *rqc2*-D98A. (See Supplementary Figures S7C-D for corresponding fluorescent polysome profiles) \*\**P* < 0.01; \**P* < 0.05; ns, not significant, Student’s *t* test. **(D)** Polysome profiles of lysates were prepared from *ltn1Δ* cells with WT *RQC2*, *rqc2*-D9A and *rqc2*-D98A after 8 hours of cycloheximide treatment. **(E-F)** Areas under the 60S, 80S and polysome peaks (up to tetrasomes) from the sucrose gradient profiles were quantified for each strain across three independent experiments. The ratio of areas covered by 60S and 80S peaks was plotted in **(E)**; ratio of polysomes to 80S was plotted in **(F)**. \*\**P* < 0.01; \**P* < 0.05; Student’s *t* test. **(G)** Equivalent number of yeast cells were spotted in 10-fold serial dilutions on YPD plates, with or without cycloheximide, and grown for 60 hrs before imaging. **(H)** Model for modes of CAT tailing and dysfunction. *Left*, in ‘extrusion mode’, lack of pulling force facilitates P-site tRNA (in pink) mobility and allows for incorporation of both Ala and Thr into CAT tails to extrude stalled NCs from the 60S exit tunnel into the extra-ribosomal space. Thr disfavors the formation of polyalanine α-helices that interfere with extrusion. NC extrusion increases the accessibility of NC lysines to Ltn1p and thus promotes ubiquitylation. *Top right*, after NC ubiquitylation, pulling by Cdc48p transmits a force along the NC that limits P-site tRNA mobility and switches CAT tailing to ‘release mode’. This leads to the addition of Ala-only CAT tails that aid in the release of the NC. Ala CAT tails act as degrons off the ribosome. *Bottom right*, the absence of pulling force leads to accumulation of RNP complexes and NCs with long, Thr-rich tails. These tails are inefficiently released, prone to aggregation, and function poorly as degrons. Ala-tRNA is shown in teal, Thr-tRNA in gold.

The CAT tail deficient strain (*rqc2*-D98A) has a growth defect under CHX-induced stalling stress when LTN1 is deleted^22^. To assess whether defects in NC release may contribute to this growth phenotype, we assessed growth and ribosome state in these conditions using spot assays and polysome profiling, respectively. In the presence of CHX, slow growth and abnormal polysome traces were observed when CAT tailing was disabled in rqc2-D98A (Figure 7E, 7H). The ratio of 60S/80S increased and polysomes/80S decreased (Figure 7F-G). This is consistent with increased 60-NC and 80S-NC (recombined *in vitro*) complexes that cannot be used for translation. The *rqc2*-D9A strain had no defect in polysomes or growth (Figure 7E, 7G), suggesting that short CAT tails may be sufficient for release. An intermediate level of release was measured for our model substrates in Rqc2-D9A (Figure 7C). Together, our data argue that pulling force on the NC induces short Ala CAT tailing that promotes nascent chain release from 60S ribosomal subunits and appends a degron to the NC. We term this form of CAT tailing “release mode”

## DISCUSSION

### Mechanochemical forces on the NC dictate CAT Tail composition and function

Our findings support a model in which mechanochemical forces dictate the composition of CAT tails and this determines their function and dysfunction (Figure 7H). Before the NC is ubiquitylated, CAT tailing operates in an “extrusion mode” that incorporates both Ala and Thr into the stalled NC and retains the NC on the 60S ribosomal subunit. These Ala/Thr CAT tails function to extrude the NC from the 60S and increase accessibility of lysines on the NC for Ltn1p to ubiquitylate. The presence of Thr in CAT tails restricts the formation of polyalanine α-helices and this increases the efficiency of NC extrusion (Figure 6). After NC ubiquitylation, the AAA+ ATPase Cdc48 generates a pulling force that switches CAT tailing to “release mode”. In release mode, short Ala CAT tails are added to the NC that promote release of NC from the 60S ribosomal subunit. This terminates CAT tailing, recycles the 60S ribosomal unit, and leaves the NC with a C-terminal CAT tail degron that functions in parallel to ubiquitin to target the NC for destruction. In the absence of mechanochemical forces that transition CAT tailing from extrusion mode to release mode, a buildup of “clogged” 60S-NC ribosome subunits occurs that can impair translation (Figure 7). The presence of Thr in CAT tails increases the efficiency of pulling forces in switching CAT tail to extrusion to release mode, likely through restricting the formation of polyalanine α-helices. Off the ribosome, Thr-rich CAT tails aggregate and induce proteotoxic stress. Mechanochemical forces can originate from multiple sources, most prominently pulling by Cdc48p (Figure 4), but also extra-ribosomal NC folding (Figure 3), NC interactions with a membrane translocon (Figure 4), electrostatic interactions between the NC and ribosomal exit tunnel (Figure 5), and potentially, NC interactions with chaperones^79^.

### CAT Tail composition selectivity

We propose two non-mutually exclusive mechanistic models to explain how pulling forces lead to distinct CAT tail compositions (Figure S7E). In the ‘energy barrier model’, mechanochemical forces on the NC constrict the position and entropy of the P-site tRNA, which in turn differentially affect the incorporation of Thr or Ala into the CAT tail. This is plausible because mechanical forces on the NC have been demonstrated to be potent regulators of P-site tRNA position^69,80^, in this way acting as regulators of canonical translation^81–83^. Yet unlike canonical translation, where tRNA selection is primarily driven by codon-anticodon interactions, we propose that the state of the P-site tRNA may influence the incorporation of specific amino acids during CAT tailing. Comparison of peptide bond formation energetics suggests that the electron-withdrawing nature of the Thr side chain hydroxyl can reduce the nucleophilicity of the ⍺-amino group, thereby elevating the activation energy required for peptide bond formation^84^. In the absence of mechanochemical pulling forces, the higher entropy and degrees of freedom of the P-site tRNA may help overcome such an energetic barrier, facilitating the incorporation of both Thr and Ala in CAT tails. Conversely, when pulling forces limit P-site tRNA mobility, Ala incorporation is favored due to a lower energy threshold for peptide bond formation. The ‘tRNA recruitment model’ proposes that mechanochemical forces on the P-site tRNA allosterically modulate the recruitment of charged tRNAs at the A-site via Rqc2p interactions. Structural analysis of the NC-60S-Rqc2p complex reveals that the P-site tRNA is coordinated by both NFACT-R/C and NFACT-N domains of Rqc2p, while the A-site tRNA primarily contacts the NFACT-N domain. These domains are linked by a coiled-coiled middle (CC-M) domain. This model suggests that pulling forces, which reduce P-site tRNA mobility, alter the NFACT-N domain’s affinity for Thr-tRNA, thereby favoring Ala-tRNA recruitment and incorporation during CAT tailing. Structural studies may reveal if Rqc2p makes additional contacts in the release mode that would favor recruitment of Ala tRNAs. Future biochemical, structural, and *in silico* studies can distinguish the contributions of these non-mutually exclusive models.

### NC release from the 60S ribosomal subunit

We here propose that Ala CAT tailing leads to efficient NC release from 60S ribosomal subunits. Studies in bacteria and humans have proposed that non-ubiquitinated NCs have their NC-tRNA ester bond cleaved by dedicated peptidyl-tRNA hydrolases, such as Pth (bacteria)^29^ and Ptrh1 (human)^28^. Our findings that Ala CAT tails favor NC release are consistent with a recent study in bacteria that proposed that Ala-tails are preferred substrates for the Pth enzyme that cleaves the NC-tRNA bond^29^. This work argues that Ala tailing may serve to counteract NC-generated pulling force on the P-site tRNA and enhance NC-tRNA ester bond access for Pth. We speculate this may relate to the formation of α-helices by Ala CAT tails. Another mechanism to explain the amino acid dependance of NC release may be that the NC-tRNA bond undergoes spontaneous hydrolysis with the timing dependent on the identity of the C-terminal amino acid. This is consistent with observations that alanyl-tRNA has a shorter half-life in water than threonyl-tRNA (6 min vs 38 min)^11,85^. In this model, incorporation of Thr into a CAT tail better allows additional residues to be added before spontaneous hydrolysis occurs and thus favors longer CAT tails. By contrast, incorporation of Ala provides limited opportunities for further Ala CAT tailing due to the relatively short half life of alanyl-tRNA and thus leads to shorter CAT tails. Water may also be specifically positioned to rapidly drive tRNA hydrolysis and NC release of Ala CAT tails. NCs are also proposed to undergo tRNA cleavage mediated by Vms1 (yeast)^26,27^ and ANKZF1 (human)^28,48^ when the NC is ubiquitylated. However, deletion of *VMS1* had no effect on the CAT tail length phenotypes in both wild-type and *ltn1Δ* backgrounds. Therefore, either Vms1p is not involved in CAT tailing and release of our substrates or it acts post-CAT tailing. We also note that our data suggests that NC pulling may reduce the affinity of Rqc2p for the 60S-NC (allowing reassociation with 40S *in vitro*) and that this may also affect NC release. Further work will help to distinguish between models of CAT tail termination and NC release.

Our results indicate that the maximum length of CAT tails observed for our substrates was approximately 70 amino acids, suggesting that this may be an upper size limit to CAT tailing. We speculate that this limit could be imposed by the interaction of chaperones with the CAT-tailed NC or, alternatively, by the formation of globular structures within the CAT tail itself. Both of these could generate mechanical pulling forces on the NC that terminate CAT tailing. This size limitation could be beneficial in preventing excessively long CAT tails and ensuring that every CAT tailing event ends in release mode, allowing the nascent chain to be released from the ribosome even in the absence of ubiquitination and protein folding.

### Evolutionary implications of CAT tailing in eukaryotes

We propose that the evolution of CAT tailing from prokaryotes to eukaryotes included the addition of Thr residues to disrupt polyalanine α-helices and thereby efficiently extend the NC on the 60S ribosomal subunit to increase accessibility of the NC to Ltn1p, which is not present in bacteria. Our data also argues that force transfer to the PTC via nascent chain pulling by Cdc48p is inefficient if polyalanine α-helices are present, and that Thr in CAT tails thus facilitates force transfer along that nascent chain which is critical for switching CAT tailing from extrusion to release mode. Recent studies in human neurons suggest that human CAT tails may be up to 20 amino acids in length and include amino acids such as glycine (Gly), lysine (Lys), and aspartate (Asp), in addition to Ala and Thr^15^. Mass spectrometry analysis of CAT tailed C-130 protein from flies suggests that serine (Ser) and cysteine (Cys) may also be part of CAT tails^14^. These amino acids may impart as-yet-unknown functions to CAT tails, but also likely function like Thr in preventing the formation of α-helices within the ribosome exit tunnel as would occur with polyalanine. This allows pulling force on the NC to efficiently switch CAT tailing from ‘extrusion mode’ to ‘release mode’. However, unlike Ala tails, Ala/Thr tails do not promote the efficient release of ubiquitylated NCs from the 60S subunit and are prone to aggregation. Thus, the evolution of Thr in CAT tails represents a trade-off between promoting Ltn1p-mediated ubiquitination on the ribosome and the potential for inefficient NC release from ribosomes and CAT aggregation. How the consequences of these trade-offs may contribute to neurodegenerative disease is an attractive area for future study.

The mechanistic principles underlying CAT tailing may provide a window into the primordial origins of protein synthesis. The RNA world hypothesis proposes that early protein synthesis evolved from simple RNA-based systems. Notably, peptide bond formation in both CAT tailing and canonical translation can be explained by reaction energetics, P-site tRNA distortion, and tRNA translocation thermodynamics, without requiring GTP hydrolysis^3,86–88^. Our results show that NC/exit tunnel and NC/extra-ribosomal factor interactions are sufficient to specify the sequence of CAT tails. This suggests that in the primordial world, physical interactions between nascent primitive peptides and their environment could have allowed for the production of a diverse and stereotyped peptidome. Such peptides may have significantly expanded the structural and functional repertoire of early biomolecules, potentially playing a crucial role in the transition from an RNA world to the protein-rich life we observe today

### Natural RQC substrates and disease

Our results provide a framework for understanding the behavior of natural RQC substrates. Due to readthrough into the poly(A) tail, NCs from non-stop mRNAs often contain poly-Lys tracts positioned within the ribosomal exit tunnel. Based on our findings, we speculate that the positive charges of the poly-Lys tract may generate pulling forces that favor the formation of short, Ala-rich CAT tails, promoting efficient release and off-ribosome degradation of these aberrant NCs but limited extrusion of the NC. Another category of RQC substrates are mRNAs translated at ER^89^ or mitochondrial membranes^14,47^. These proteins are predicted to receive short Ala tails due to mechanochemical forces that occur during cotranslational insertion at the ER and mitochondria. Indeed, short CAT tails were observed on ER-RQC substrates in human cells^16^. However, defects in cotranslational insertion may decrease these forces, increasing the frequency of Thr CAT tailing^89^. Moreover, in such cases proteins are likely to be misfolded due to improper retention in the cytosol, which reduces intrinsic pulling forces and further favors the addition of Thr in CAT tails. Consistent with this, CAT tailing of membrane proteins with defective cotranslational insertion into mitochondria^14^ and ER^39^ has been proposed to drive cytotoxic aggregation and disease phenotypes. The etiological role of dysfunctional CAT tailing in the context of human disease as well as potential therapeutic implications of modulating this pathway are promising areas of investigation.

### Limitations of the study

Our study does not precisely determine the primary sequence and length of CAT-tailed proteins formed under different pulling forces due to technical challenges associated with sequencing a heterogeneous ensemble of proteins. Instead, we perform bulk analysis of differing substrates, including amino acid analysis and western blots. Our study does not definitively explain how pulling forces bias the incorporation of Ala or Thr but instead offers orthogonal experiments supporting this phenomenon and suggests potential mechanisms in the discussion section. For the polysome profiling experiments in low salt, we interpret the population of 60S which does not associate with 40S into 80S as having steric obstruction at the 40S/60S interface. We hypothesize that the non-associating 60S-RQC substrate population we observe is bound to RQC factors (e.g. Rqc2p or Ltn1p), however other factors may influence this as well, including the amount of 40S that are capable of associating with 60S. These caveats should be considered when interpreting our results.

## MATERIALS AND METHODS

### Yeast strains and growth conditions

All plasmids (including DNA and encoded amino acid sequences) and yeast strains used in this study are listed in Supplementary Table S1. All yeast cultures were grown at 30°C in synthetic complete (SC) media supplemented with appropriate nutrient dropouts. Deletion strains were generated primarily in the BY4741 background through transformation with PCR products containing antibiotic selection cassettes generated with Phusion polymerase (New England Biolabs). All selection markers were integrated using homologous integration. Experiments involving *cdc48-3* mutants and *LTN1/RQC2* deletion were performed in the W303 background. Successful transformants were confirmed by PCR using SapphireAmp Fast PCR Master Mix.

### Plasmid construction

All plasmids were derived from the pRS316 vector and constructed using the Gibson assembly (New England Biolabs HiFi DNA Assembly Master Mix). The assembled plasmids were then transformed into competent *E. coli* DH5α cells using the heat shock method. Transformed cells were plated on Luria-Bertani (LB) agar plates containing 100 μg/ml carbenicillin and incubated overnight at 37°C. Single colonies of *E. coli* transformants were picked and grown overnight in 5 ml LB medium supplemented with 100 μg/ml carbenicillin at 37°C. Plasmids were extracted from these overnight cultures using the homemade miniprep kit and subsequently validated by Sanger sequencing (McLab) or NGS sequencing (Plasmidsaurus).

### Immunoblots

For whole-cell immunoblot analysis, log-phase yeast cultures derived from a single colony were harvested at an OD600 of 0.8 by centrifugation. In experiments involving AZC, canavanine and bortezomib treatments, yeast cultures at OD600 of 0.5 were exposed to the relevant drugs for a duration of 3 hours. The volume of culture collected was adjusted to yield a cell density equivalent to 0.25 OD600 units per milliliter. The cell pellets were then resuspended in 30 μl of 4× NuPage LDS Sample Buffer containing 5% β-mercaptoethanol. To facilitate cell lysis and protein denaturation, the resuspended pellets were boiled at 95°C for 5 minutes. SDS-PAGE was performed using Novex NuPAGE 4–12% Bis-Tris gels (Thermo Fisher Scientific) in all cases except where otherwise stated. The separated proteins were then transferred onto 0.45-μm nitrocellulose membranes (Bio-Rad) using a Transblot Turbo system (Bio-Rad). To prevent non-specific antibody binding, the membranes were incubated in a blocking solution, 5% fat-free milk in Tris-buffered saline with Tween 20 (TBST), for 15 minutes at room temperature. Primary antibody incubations were carried out either overnight at 4°C or for 4 hours at room temperature. The following primary antibodies used: mouse anti-GFP (MA5-15256; Thermo Fisher Scientific), rabbit anti-Hexokinase (H2035-01; US Biological), mouse anti-MBP (E8032S; NEB) and Pierce mouse anti-HA (26183; Invitrogen). After primary antibody incubation, the blots were washed 3 times in TBST before being probed with appropriate secondary antibodies: IRDye 800CW donkey anti-mouse, IRDye 680RD goat anti-rabbit, or IRDye 680RD goat anti-mouse (LiCor Biosciences). All antibodies were used at a 1:5000 dilution. The immunoblots were visualized and digitally captured using a LiCor Odyssey imaging system. All original uncropped blots for immunoblots are included in Supplemental Document S1.

### CAT tail length and aggregation analyses

Densitometric analyses of western blots were performed with raw images using Image Studio Lite software. A box of fixed dimensions was drawn around the well region to quantify aggregated proteins and expressed as a percentage of reporter with maximum aggregation (See Supplemental Document S2 for details). To calculate CAT tail lengths, a semi-log plot of molecular weights of ladder was generated and from the resulting best fit plot, the maximum molecular weight shift between CAT tailed (*ltn1Δ RQC2*) and non-CAT tailed (*ltn1Δ rqc2Δ*) species was calculated. Assuming the average amino acid mass as 110 Da, CAT tail length was estimated (See Supplemental Document S2 for details).

### Immunoprecipitation and amino acid analysis

Yeast transformants were grown to around OD600=0.6 and harvested by vacuum filtration, flash-frozen in liquid nitrogen and stored at −80°C until use. Cells were lysed by cryo-grinding in a Spex 6750 Freezer Mill in two 10 Hz runs of 2 mins, each followed by a gap of 1 min. Pulverized cells were solubilized in the immmunoprecipitation (IP) buffer consisting of 50 mM HEPES pH 7.4, 100 mM NaCl and Roche cOmplete Protease inhibitors. The resulting lysate was clarified by centrifugation at 5,000g for 5 min followed by two rounds of centrifugation at 22,000g for 15 min. GFP was isolated from the clarified lysate using magnetic agarose GFP-Trap (Chromotek) resin as bait. After six washes with the IP buffer, the bound GFP was eluted with a buffer consisting of 50 mM HEPES pH 2.5,100 mM NaCl. Amino acid analysis was performed with eluted proteins at the UC Davis Genome Center.

### Flow Cytometry

Flow cytometry was performed on log-phase yeast cultured in synthetic defined media using a BD Accuri C6 flow cytometer (BD Biosciences). For each condition, at least three biological replicates, derived from independent cultures of separate colonies, were analyzed. A minimum of 10,000 events were collected for each sample. The acquired data were processed using MATLAB as previously described^22^. Briefly, to isolate the population of yeast expressing the plasmid of interest, gating was performed based on the red fluorescent protein (RFP) fluorescence signal in plasmids harboring tandem T2A sequences separating RFP and GFP. MATLAB 23.2.0 (Mathworks) was used to calculate RFP-normalized GFP levels and log2 of mean scores calculated. Mean values based on linear scale were plotted using Prism 10.2.2 (Graphpad). Error bars represent standard deviations between biological replicates.

### Yeast Spot Assay

Log-phase yeast cultures were diluted to OD600 of 0.6 and spotted in 10-fold dilution series onto YPD plates supplemented with vehicle control DMSO or indicated concentrations of cycloheximide. Plates were incubated for 72 h for at the indicated temperatures before imaging.

### Fluorescent Polysome Profiling

Yeast transformants harboring reporter plasmids were cultured to an OD600 of 0.6. These cells were then harvested by vacuum filtration and immediately flash-frozen in liquid nitrogen. For polysome profiling experiments under cycloheximide stress, yeast strains were first grown to an OD600 of 0.4 and then 125 μg/mL cycloheximide was added to the culture. The cells were then allowed to grow for an additional 8 hours before being harvested. For fluorescent polysome profiling under ‘low salt’ experiments the lysis buffer was prepared with 20 mM Tris-HCl (pH 7.5), 140 mM KCl, 15 mM MgCl2, 100 µg/mL cycloheximide, 1% Triton X-100, 1 mM DTT, cOmplete protease inhibitor cocktail tablets (EDTA-free), and RNasin inhibitor (Promega). For experiments done under ‘high salt’ conditions, the KCl concentration in the lysis buffer was increased to 500 mM. In all cases pellets of lysis buffer were prepared in liquid nitrogen and cryo-ground with the frozen cells using a Spex 6750 Freezer Mill, once at 10 Hz for 1 minute. The lysate was then centrifuged at 1,500g for 2 minutes and the supernatant collected for immediate use in polysome profiling. A 10 - 50% sucrose gradient was prepared in Seaton Open Top Polyclear centrifuge tubes using the lysis buffer but without Triton X-100. Gradient was generated using the Gradient Master (BioComp Instruments) using manufacturer’s instructions. Equivalent amounts of cell lysates prepared as described above were loaded onto the sucrose gradients and spun at 35,000 *g* for 3 hours at 4°C in a SW41 rotor (Beckman Coulter). After centrifugation the polysomes were fractionated and absorbance was monitored at 254 nm for RNA and 680 nm for GFP using the TriaxTM Flowcell (BioComp Instruments). Areas under the curves outlined by 60S, 80S and polysome peaks were calculated by integrating the area between the curve and a line connecting the troughs on either side of each peak. This method was applied for quantifying areas under curves in the polysome profiles.

### Imaging

Yeast cells expressing RQC substrates were cultured in SD media supplemented with the necessary auxotrophic nutrients and harvested by centrifugation and resuspended in a smaller volume of SD media to increase cell density. Glass slides were coated with concanavalin A to facilitate cell immobilization. A small volume of the concentrated cell suspension was then applied to the treated slides and images were taken.

### Material availability

All DNA reporters and yeast strains used in this study are available upon request to the corresponding authors and where applicable, upon signing Material Transfer Agreements with Stanford University.

## Acknowledgments

We thank D. Herschlag, R. Kopito, B. Lu, P. Harbury, F. Scavone, and A. Frost for discussions and helpful comments in the preparation of this manuscript. We also thank members of Brandman, Kopito and Lu Labs for helpful discussions and support. This work was supported by the NIGMS of the National Institutes of Health Grant no. R35GM153301 to O.B. D.K was supported by the Stanford Dean’s Fellowship (Bernard Cohen Postdoctoral Fellowship Fund) and the Mikitani Cancer Fellowship Award of the Stanford Cancer Institute. The content is solely the responsibility of the authors and does not necessarily represent the official views of the National Institutes of Health or Stanford University. The authors declare no competing interests.

## Author contributions

Conceptualization, D.K., A.A.V., C.S.S. and O.B.; Methodology, D.K., A.A.V., C.S.S. and O.B.; Validation, D.K. and A.A.V.; Formal Analysis, D.K. and O.B.; Investigation, D.K. and A.A.V.; Resources, O.B.; Data Curation, D.K. and O.B.; Writing – Original Draft, D.K. and O.B.; Writing – Review and Editing D.K., A.A.V., C.S.S. and O.B.; Visualization, D.K.; Supervision, O.B.; Funding Acquisition, O.B. and D.K.

## Supplemental information

Document S1: Original uncropped blots for Western Blot images.

Document S2: Densitometry analysis of aggregation, CAT tail length estimation and ratio of long to short CAT tails

Table S1. List of plasmids and yeast strains used in the study, and full amino acid and DNA sequences of RQC reporters

---

**Figure S1.**
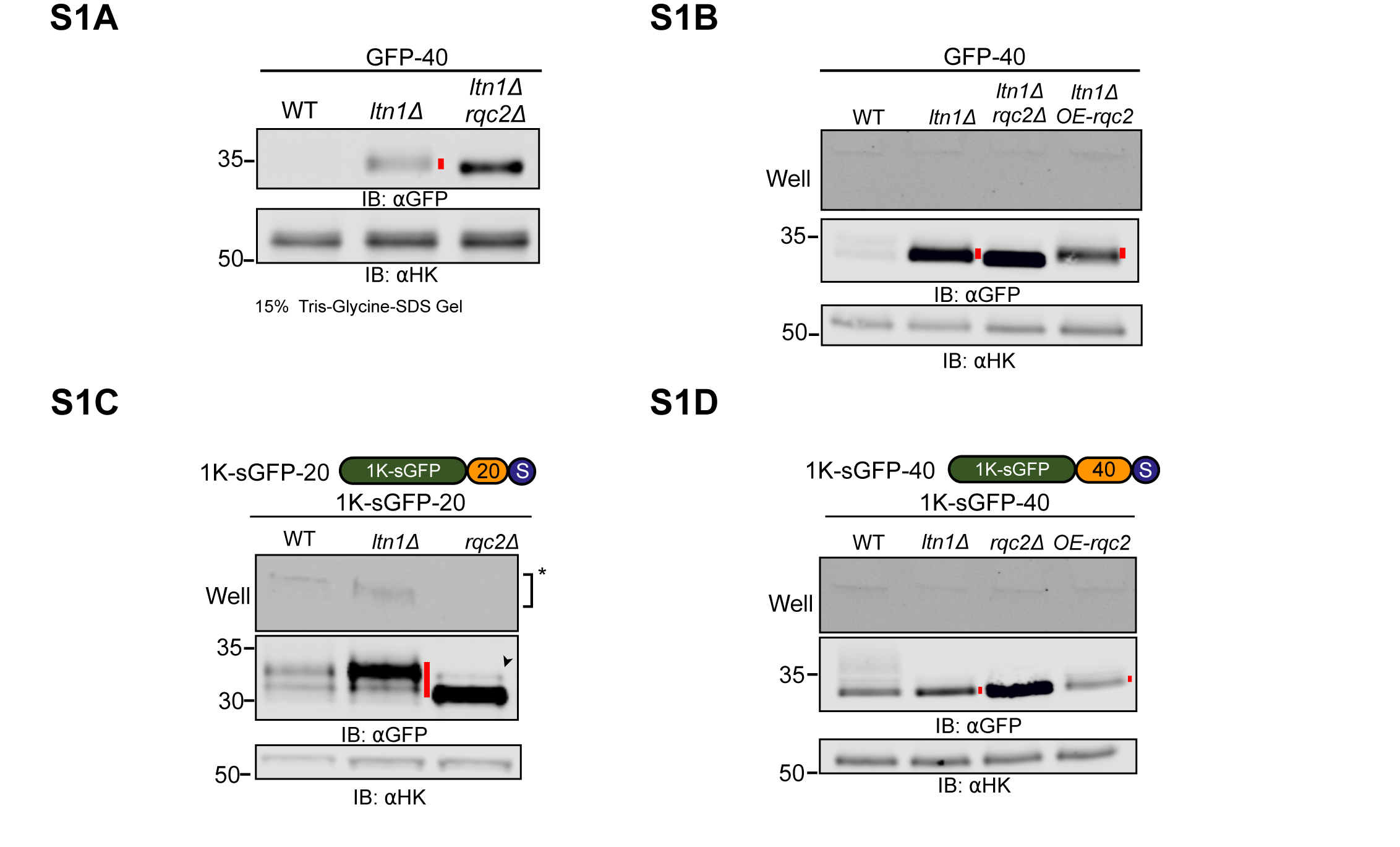
CAT tails have diverse composition, degradation, and aggregation propensities. **(A-D)** Whole cell IBs of lysates containing model RQC substrates expressed in different strains and gel densities. Cartoon of model RQC substrates with single lysine superfolder GFP (1K-sGFP) indicated where applicable. CAT tails are marked by a red line. Arrowhead (➤) indicates readthrough product formed after bypassing the polyarginine arrest sequence.

**Figure S2.**
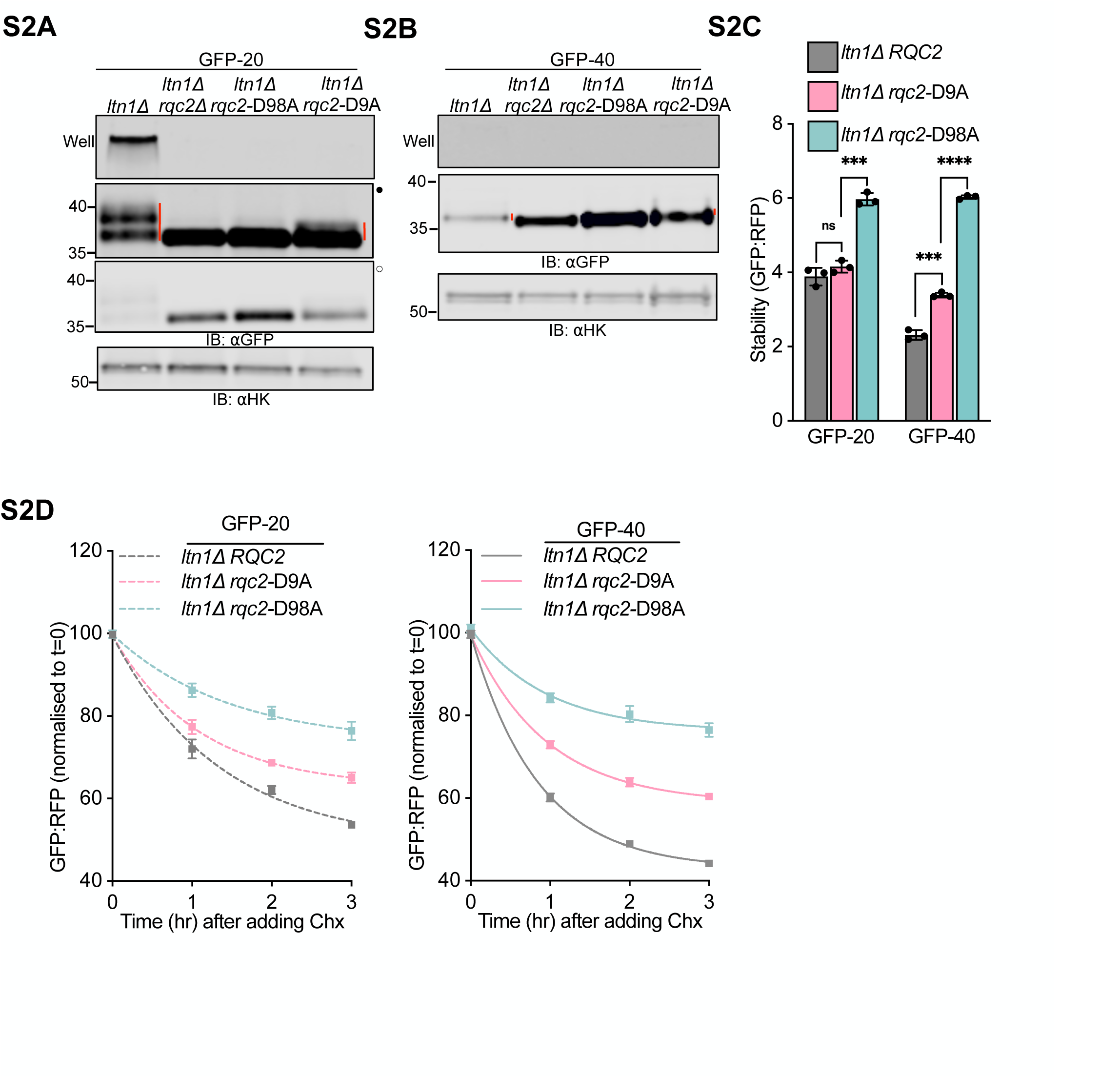
CAT tail sequence determines degradation and aggregation. **(A-B)** Whole cell immunoblots of GFP-20 and GFP-40 expressed in *ltn1Δ* cells with varying CAT tailing capabilities: normal CAT tailing (WT, lane 1), no CAT tailing (*rqc2Δ,* lane 2; *rqc2*-D98A, lane 3), and impaired CAT tailing (*rqc2*-D9A, lane 4). **(C)** Mean stability measurements for GFP-20 and GFP-40 substrates in cells with normal, impaired and no CAT tailing. \*\*\*\**P* < 0.0001; \*\*\**P* <0.001; ns, not significant. **(D)** Normalized stability measurements for GFP-20 and GFP-40 reporters expressed in *ltn1Δ* cells with variants of *RQC2* chased over 3 hours after introduction of 200 μg/mL cycloheximide.

**Figure S3.**
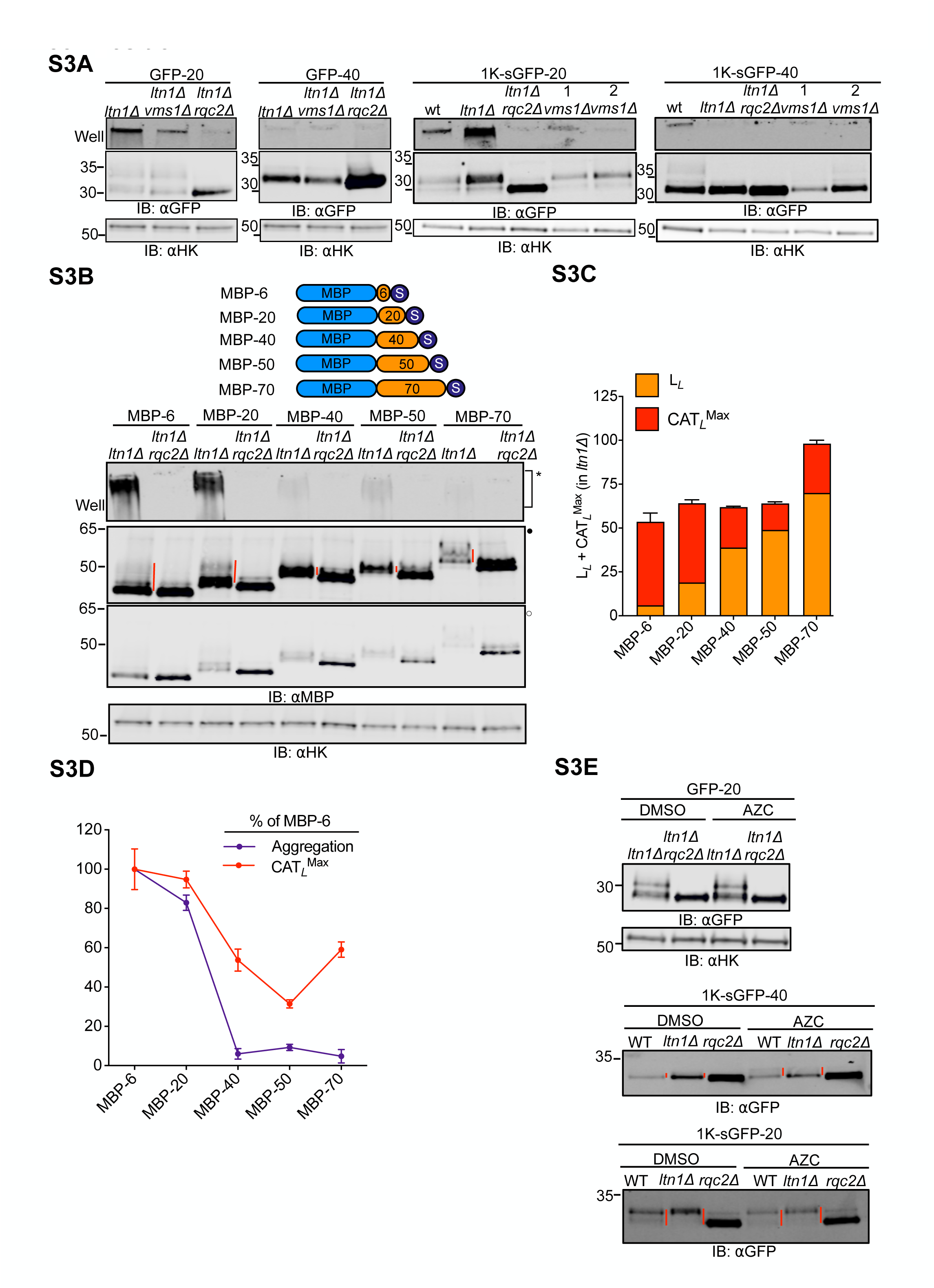
Folding-induced mechanical forces on the NC regulate CAT tail composition. **(A)** IBs with model RQC substrates expressed in different genetic backgrounds, including deletion in *VMS1*. The numbers 1 and 2 above *vms1Δ* denote two independent clones that were used in the experiments. **(B)** Cartoon of maltose binding protein (MBP) based RQC substrates with linker of varying lengths (0, 20, 40, 50 and 70 amino acids) between MBP and stalling sequence, analyzed by immunoblotting. **(C)** Stacked bar graph showing the sum of linker and estimated maximum CAT tail length for each RQC substrate in the *ltn1Δ* background. **(D)** Relative aggregation and maximum CAT tail length for each RQC substrate, normalized to the MBP-6 substrate, in the *ltn1Δ* background. **(E)** Cells expressing GFP-20, 1K-sGFP-40 and 1K-sGFP-20 were grown in media supplemented with DMSO or 1 mM AZC for 3 hours and analyzed by immunoblotting.

**Figure S4.**
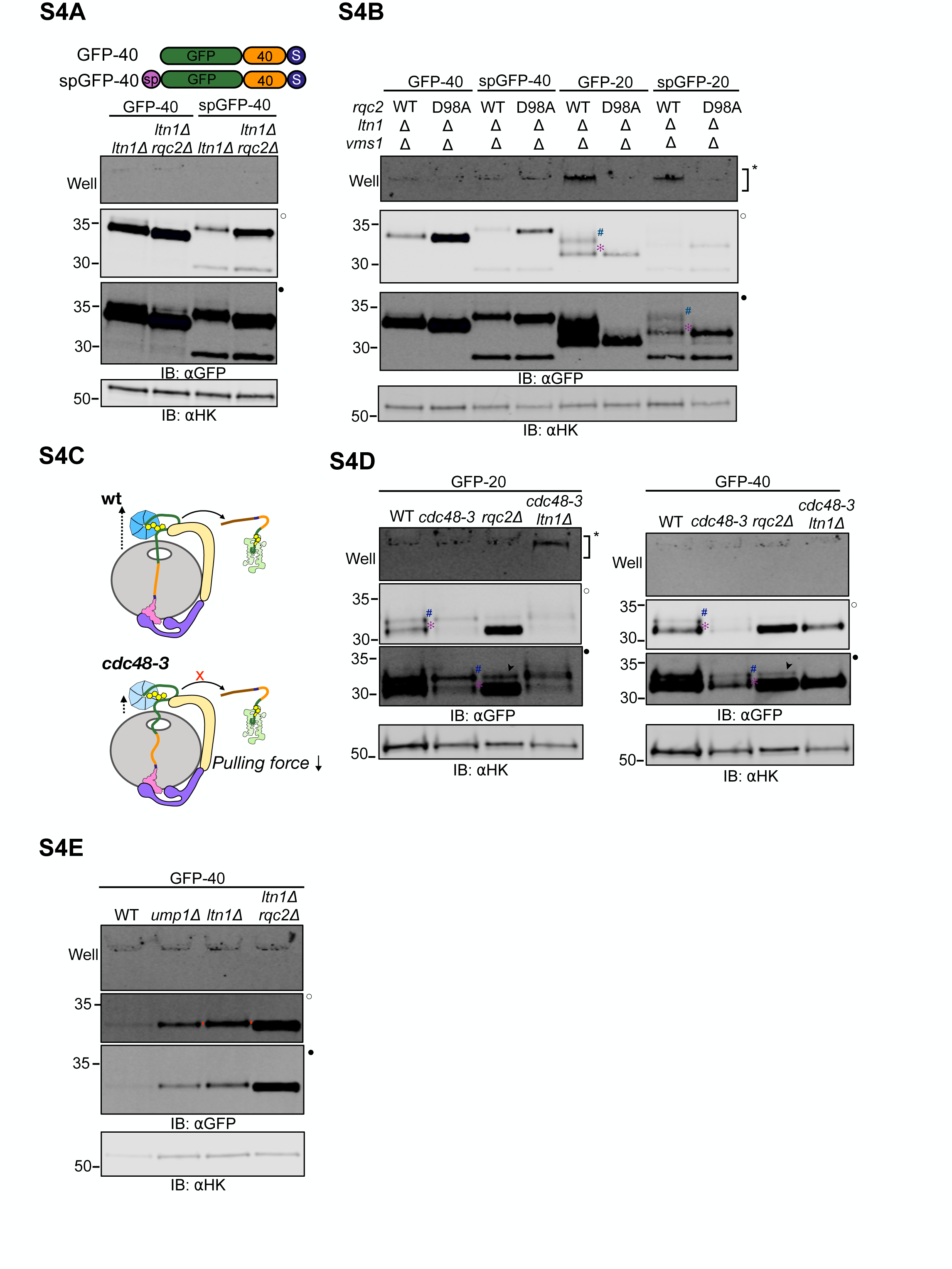
Extrinsic mechanical forces on the NC determine CAT tail sequence. **(A)** Cartoon of ER-directed RQC substrate spGFP-40 alongside the cytosolic variant, with the IBs. **(B)** IBs with model cytosolic and ER directed RQC substrates expressed in *ltn1Δ vms1Δ* background with WT and mutant *RQC2*. **(C)** Schematics showing the RQC role of *CDC48* in different backgrounds. *Top*, in WT cells Cdc48p binds and extracts the ubiquitylated NC from the 60S for proteasomal degradation. *Bottom*, the Cdc48p activity is reduced in the *cdc48-3* background thereby decreasing the pulling force exerted on the NC. **(D)** GFP-20 and GFP-40 reporters expressed in cells with WT and mutant *CDC48*, analyzed by immunoblotting along with the no-CAT tailing (*rqc2Δ*) and no ubiquitylation (*cdc48-3 ltn1Δ*) controls. Long and short CAT tails are marked by hash (#) and asterisk (✻) respectively. Arrowhead (➤) indicates readthrough product formed after bypassing the polyarginine arrest sequence. **(E)** GFP-40 reporter expressed in *ump1Δ* cells, compared with WT, non-ubiquitylated (*ltn1Δ*), and non-CAT-tailed (*rqc2Δ*) controls

**Figure S7.**
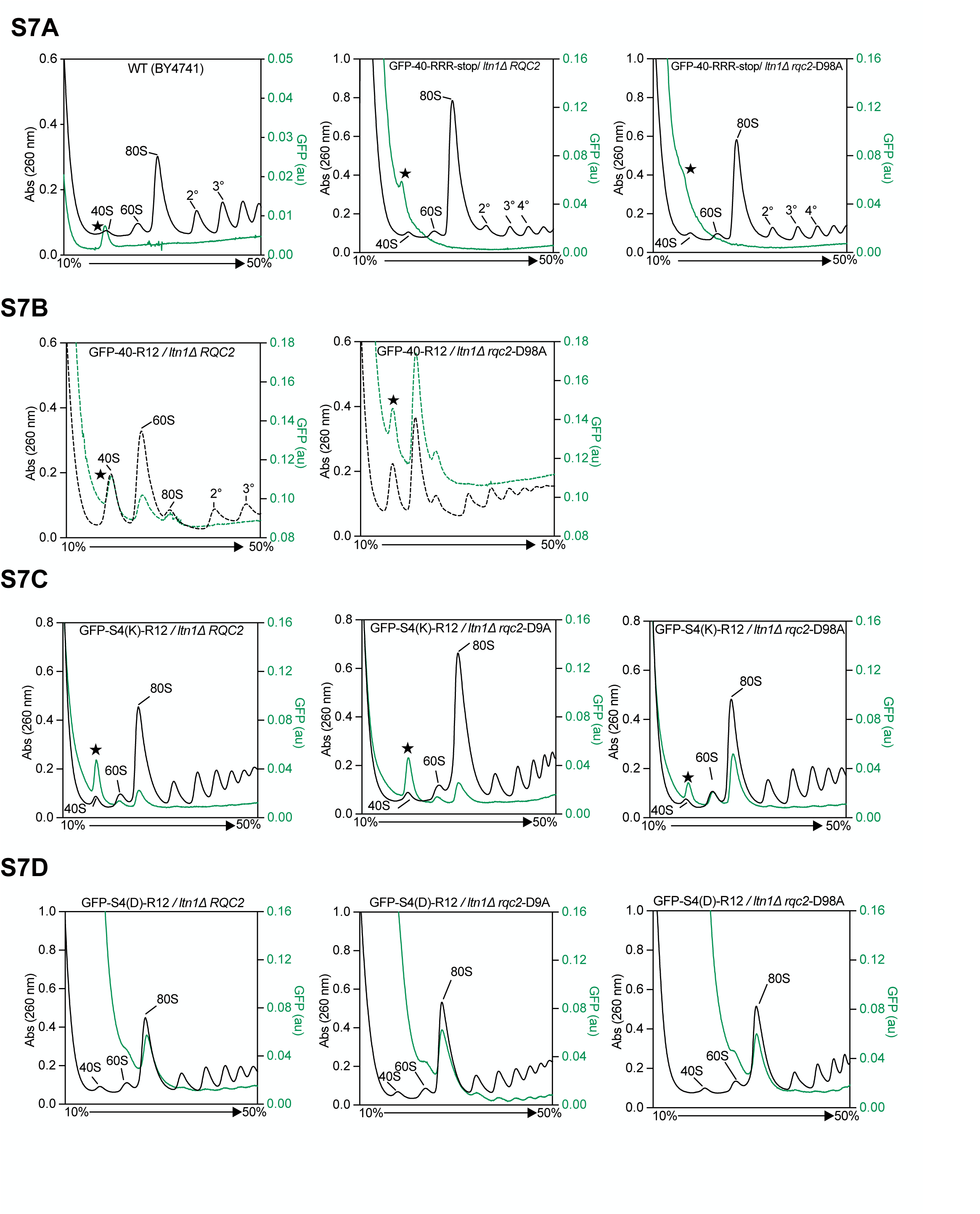
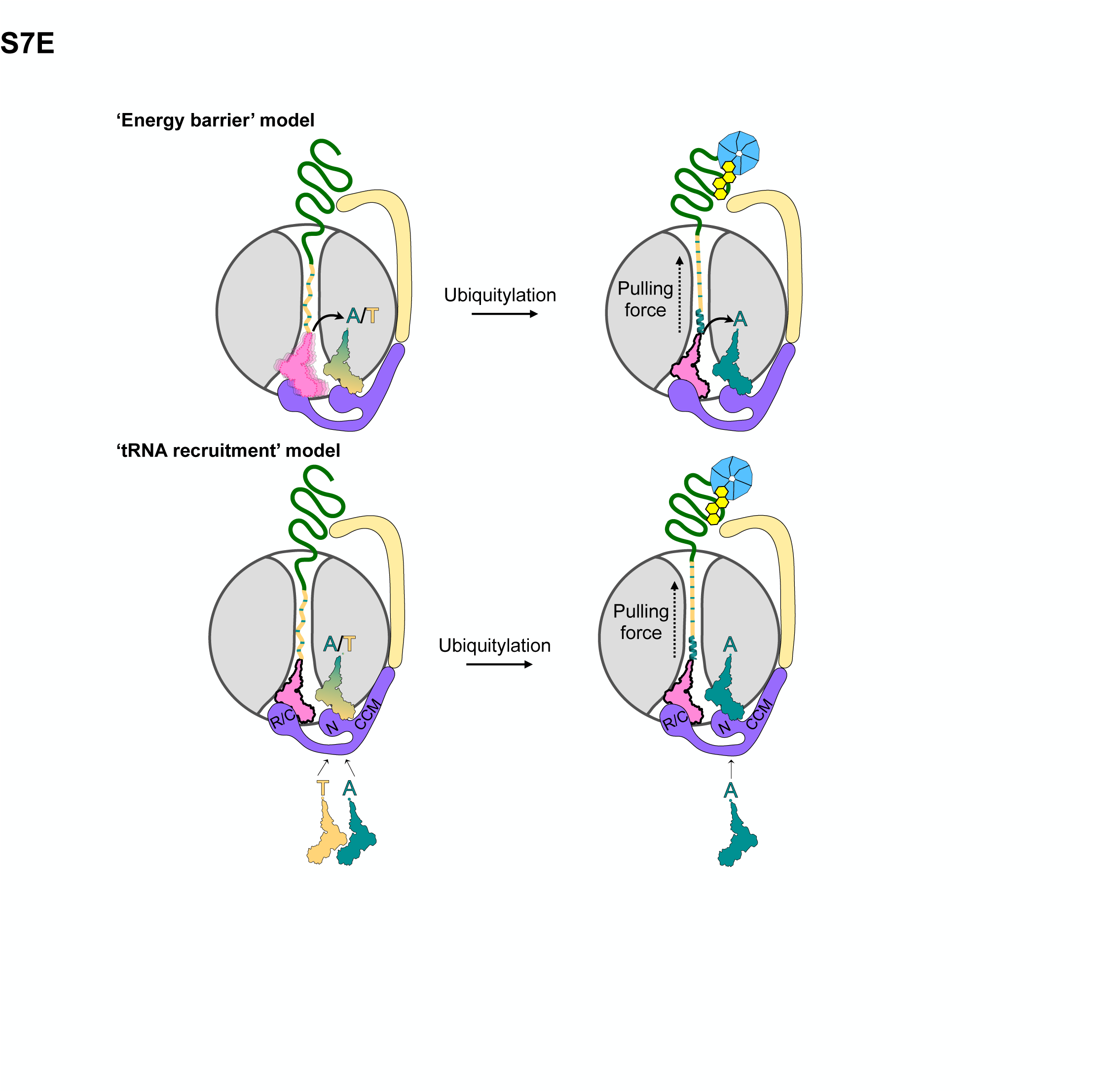
CAT tails regulate NC retention and release. **(A)** *Left,* **Fluorescent polysome profiling of** WT cells after 8 hours of cycloheximide treatment, harboring no plasmid, showed a background fluorescence peak at 40S position; Fluorescent polysome profiling of *ltn1Δ* (*center*) and *ltn1Δ rqc2*-D98A (*right*) cells expressing GFP-40-RRR-stop plasmid as a control. **(B)** Fluorescent polysome profiling of GFP-40/*ltn1Δ* (*left*) and GFP-40/*ltn1Δrqc2-*D98A (*right*) in a ‘high salt’ buffer containing 500 mM KCl. Dotted lines in the traces indicate that profiles were collected under high salt conditions. **(C)** GFP-S4(K) and **(D)** GFP-S4(D) reporters expressed in *ltn1Δ* cells with WT *RQC2*, *rqc2*-D9A and *rqc2*-D98A, were analyzed by fluorescent polysome profiling. **(E)** Schematic representation of models for amino acid specificity in CAT tailing. *Top*, *‘Energy Barrier’* model. In the absence of a pulling force on the NC, the P-site tRNA has higher degrees of freedom, allowing it to incorporate both Ala and Thr into the CAT tail. This increased entropy of the P-site tRNA helps overcome the higher activation energy barrier for Thr incorporation. When a pulling force is applied to the NC, it restricts the P-site tRNA’s mobility and lowers its entropy. Under these conditions, Ala is selectively incorporated due to its lower activation energy barrier for peptide bond formation, allowing it to more easily accommodate the restricted state of the peptidyl transferase center. *Bottom,* ‘*tRNA recruitment* model’. Pulling forces regulate the recruitment of charged tRNAs at the A-site via allosteric interactions with Rqc2p. The NFACT-N/R and NFACT-C domains stabilize the anticodon arm of P-site tRNA, whereas the NFACT-C domain and 60S facing side of CCM loop stabilize the anticodon arm of A-site tRNAs. The NFACT-N domain specifically recruits Ala-tRNA^AGC^ and Thr-tRNA^AGT^ and delivers them to the 60S A-site. In the absence of a pulling force, recruitment of both Ala- and Thr-tRNA is favored. The presence of pulling force reduces the affinity of NFACT-N domain for Thr-tRNA, thereby favoring the recruitment of Ala-tRNA.

## Notes

### Competing Interest Statement

The authors have declared no competing interest.

### Summary of Updates

Changed the title and included new experiments in figures 6 and 7.

## REFERENCES

1. Sitron, C.S., and Brandman, O. (2020). Detection and Degradation of Stalled Nascent Chains via Ribosome-Associated Quality Control. Annu. Rev. Biochem. 89, 417–442. 10.1146/annurev-biochem-013118-110729.

2. Yip, M.C.J., and Shao, S. (2021). Detecting and Rescuing Stalled Ribosomes. Trends Biochem. Sci. 46, 731–743. 10.1016/j.tibs.2021.03.008.

3. Howard, C.J., and Frost, A. (2021). Ribosome-associated quality control and CAT tailing. Crit. Rev. Biochem. Mol. Biol. 56, 603–620. 10.1080/10409238.2021.1938507.

4. Inada, T., and Beckmann, R. (2024). Mechanisms of Translation-coupled Quality Control. J. Mol. Biol. 436, 168496. 10.1016/j.jmb.2024.168496.

5. Brandman, O., Stewart-Ornstein, J., Wong, D., Larson, A., Williams, C.C., Li, G.-W., Zhou, S., King, D., Shen, P.S., Weibezahn, J., et al. (2012). A ribosome-bound quality control complex triggers degradation of nascent peptides and signals translation stress. Cell 151, 1042–1054. 10.1016/j.cell.2012.10.044.

6. Shao, S., Brown, A., Santhanam, B., and Hegde, R.S. (2015). Structure and assembly pathway of the ribosome quality control complex. Mol. Cell 57, 433–444. 10.1016/j.molcel.2014.12.015.

7. Defenouillère, Q., Yao, Y., Mouaikel, J., Namane, A., Galopier, A., Decourty, L., Doyen, A., Malabat, C., Saveanu, C., Jacquier, A., et al. (2013). Cdc48-associated complex bound to 60S particles is required for the clearance of aberrant translation products. Proc. Natl. Acad. Sci. U. S. A. 110, 5046–5051. 10.1073/pnas.1221724110.

8. Juszkiewicz, S., Speldewinde, S.H., Wan, L., Svejstrup, J.Q., and Hegde, R.S. (2020). The ASC-1 Complex Disassembles Collided Ribosomes. Mol. Cell 79, 603–614.e8. 10.1016/j.molcel.2020.06.006.

9. Matsuo, Y., Tesina, P., Nakajima, S., Mizuno, M., Endo, A., Buschauer, R., Cheng, J., Shounai, O., Ikeuchi, K., Saeki, Y., et al. (2020). RQT complex dissociates ribosomes collided on endogenous RQC substrate SDD1. Nat. Struct. Mol. Biol. 27, 323–332. 10.1038/s41594-020-0393-9.

10. Bengtson, M.H., and Joazeiro, C.A.P. (2010). Role of a ribosome-associated E3 ubiquitin ligase in protein quality control. Nature 467, 470–473. 10.1038/nature09371.

11. Shao, S., von der Malsburg, K., and Hegde, R.S. (2013). Listerin-dependent nascent protein ubiquitination relies on ribosome subunit dissociation. Mol. Cell 50, 637–648. 10.1016/j.molcel.2013.04.015.

12. Shen, P.S., Park, J., Qin, Y., Li, X., Parsawar, K., Larson, M.H., Cox, J., Cheng, Y., Lambowitz, A.M., Weissman, J.S., et al. (2015). Protein synthesis. Rqc2p and 60S ribosomal subunits mediate mRNA-independent elongation of nascent chains. Science 347, 75–78. 10.1126/science.1259724.

13. Lytvynenko, I., Paternoga, H., Thrun, A., Balke, A., Müller, T.A., Chiang, C.H., Nagler, K., Tsaprailis, G., Anders, S., Bischofs, I., et al. (2019). Alanine Tails Signal Proteolysis in Bacterial Ribosome-Associated Quality Control. Cell 178, 76–90.e22. 10.1016/j.cell.2019.05.002.

14. Wu, Z., Tantray, I., Lim, J., Chen, S., Li, Y., Davis, Z., Sitron, C., Dong, J., Gispert, S., Auburger, G., et al. (2019). MISTERMINATE Mechanistically Links Mitochondrial Dysfunction with Proteostasis Failure. Mol. Cell 75, 835–848.e8. 10.1016/j.molcel.2019.06.031.

15. Udagawa, T., Seki, M., Okuyama, T., Adachi, S., Natsume, T., Noguchi, T., Matsuzawa, A., and Inada, T. (2021). Failure to Degrade CAT-Tailed Proteins Disrupts Neuronal Morphogenesis and Cell Survival. Cell Rep. 34, 108599. 10.1016/j.celrep.2020.108599.

16. Scavone, F., Gumbin, S.C., Da Rosa, P.A., and Kopito, R.R. (2023). RPL26/uL24 UFMylation is essential for ribosome-associated quality control at the endoplasmic reticulum. Proc. Natl. Acad. Sci. U. S. A. 120, e2220340120. 10.1073/pnas.2220340120.

17. Osuna, B.A., Howard, C.J., Kc, S., Frost, A., and Weinberg, D.E. (2017). In vitro analysis of RQC activities provides insights into the mechanism and function of CAT tailing. Elife 6. 10.7554/eLife.27949.

18. Filbeck, S., Cerullo, F., Paternoga, H., Tsaprailis, G., Joazeiro, C.A.P., and Pfeffer, S. (2021). Mimicry of Canonical Translation Elongation Underlies Alanine Tail Synthesis in RQC. Mol. Cell 81, 104–114.e6. 10.1016/j.molcel.2020.11.001.

19. Crowe-McAuliffe, C., Takada, H., Murina, V., Polte, C., Kasvandik, S., Tenson, T., Ignatova, Z., Atkinson, G.C., Wilson, D.N., and Hauryliuk, V. (2021). Structural Basis for Bacterial Ribosome-Associated Quality Control by RqcH and RqcP. Mol. Cell 81, 115–126.e7. 10.1016/j.molcel.2020.11.002.

20. Takada, H., Crowe-McAuliffe, C., Polte, C., Sidorova, Z.Y., Murina, V., Atkinson, G.C., Konevega, A.L., Ignatova, Z., Wilson, D.N., and Hauryliuk, V. (2021). RqcH and RqcP catalyze processive poly-alanine synthesis in a reconstituted ribosome-associated quality control system. Nucleic Acids Res. 49, 8355–8369. 10.1093/nar/gkab589.

21. Tesina, P., Ebine, S., Buschauer, R., Thoms, M., Matsuo, Y., Inada, T., and Beckmann, R. (2023). Molecular basis of eIF5A-dependent CAT tailing in eukaryotic ribosome-associated quality control. Mol. Cell 83, 607–621.e4. 10.1016/j.molcel.2023.01.020.

22. Sitron, C.S., and Brandman, O. (2019). CAT tails drive degradation of stalled polypeptides on and off the ribosome. Nat. Struct. Mol. Biol. 26, 450–459. 10.1038/s41594-019-0230-1.

23. Li, S., Wu, Z., Tantray, I., Li, Y., Chen, S., Dong, J., Glynn, S., Vogel, H., Snyder, M., and Lu, B. (2020). Quality-control mechanisms targeting translationally stalled and C-terminally extended poly(GR) associated with ALS/FTD. Proc. Natl. Acad. Sci. U. S. A. 117, 25104–25115. 10.1073/pnas.2005506117.

24. Kostova, K.K., Hickey, K.L., Osuna, B.A., Hussmann, J.A., Frost, A., Weinberg, D.E., and Weissman, J.S. (2017). CAT-tailing as a fail-safe mechanism for efficient degradation of stalled nascent polypeptides. Science 357, 414–417. 10.1126/science.aam7787.

25. Shao, S., and Hegde, R.S. (2014). Reconstitution of a minimal ribosome-associated ubiquitination pathway with purified factors. Mol. Cell 55, 880–890. 10.1016/j.molcel.2014.07.006.

26. Zurita Rendón, O., Fredrickson, E.K., Howard, C.J., Van Vranken, J., Fogarty, S., Tolley, N.D., Kalia, R., Osuna, B.A., Shen, P.S., Hill, C.P., et al. (2018). Vms1p is a release factor for the ribosome-associated quality control complex. Nat. Commun. 9, 2197. 10.1038/s41467-018-04564-3.

27. Verma, R., Reichermeier, K.M., Burroughs, A.M., Oania, R.S., Reitsma, J.M., Aravind, L., and Deshaies, R.J. (2018). Vms1 and ANKZF1 peptidyl-tRNA hydrolases release nascent chains from stalled ribosomes. Nature 557, 446–451. 10.1038/s41586-018-0022-5.

28. Kuroha, K., Zinoviev, A., Hellen, C.U.T., and Pestova, T.V. (2018). Release of Ubiquitinated and Non-ubiquitinated Nascent Chains from Stalled Mammalian Ribosomal Complexes by ANKZF1 and Ptrh1. Mol. Cell 72, 286–302.e8. 10.1016/j.molcel.2018.08.022.

29. Svetlov, M.S., Dunand, C.F., Nakamoto, J.A., Atkinson, G.C., Safdari, H.A., Wilson, D.N., Vázquez-Laslop, N., and Mankin, A.S. (2024). Peptidyl-tRNA hydrolase is the nascent chain release factor in bacterial ribosome-associated quality control. Mol. Cell 84, 715–726.e5. 10.1016/j.molcel.2023.12.002.

30. Thrun, A., Garzia, A., Kigoshi-Tansho, Y., Patil, P.R., Umbaugh, C.S., Dallinger, T., Liu, J., Kreger, S., Patrizi, A., Cox, G.A., et al. (2021). Convergence of mammalian RQC and C-end rule proteolytic pathways via alanine tailing. Mol. Cell 81, 2112–2122.e7. 10.1016/j.molcel.2021.03.004.

31. Patil, P.R., Burroughs, A.M., Misra, M., Cerullo, F., Costas-Insua, C., Hung, H.-C., Dikic, I., Aravind, L., and Joazeiro, C.A.P. (2023). Mechanism and evolutionary origins of alanine-tail C-degron recognition by E3 ligases Pirh2 and CRL2-KLHDC10. Cell Rep. 42, 113100. 10.1016/j.celrep.2023.113100.

32. Wang, X., Li, Y., Yan, X., Yang, Q., Zhang, B., Zhang, Y., Yuan, X., Jiang, C., Chen, D., Liu, Q., et al. (2023). Recognition of an Ala-rich C-degron by the E3 ligase Pirh2. Nat. Commun. 14, 2474. 10.1038/s41467-023-38173-6.

33. Choe, Y.-J., Park, S.-H., Hassemer, T., Körner, R., Vincenz-Donnelly, L., Hayer-Hartl, M., and Hartl, F.U. (2016). Failure of RQC machinery causes protein aggregation and proteotoxic stress. Nature 531, 191–195. 10.1038/nature16973.

34. Yonashiro, R., Tahara, E.B., Bengtson, M.H., Khokhrina, M., Lorenz, H., Chen, K.-C., Kigoshi-Tansho, Y., Savas, J.N., Yates, J.R., Kay, S.A., et al. (2016). The Rqc2/Tae2 subunit of the ribosome-associated quality control (RQC) complex marks ribosome-stalled nascent polypeptide chains for aggregation. Elife 5, e11794. 10.7554/eLife.11794.

35. Sitron, C.S., Park, J.H., Giafaglione, J.M., and Brandman, O. (2020). Aggregation of CAT tails blocks their degradation and causes proteotoxicity in S. cerevisiae. PLoS One 15, e0227841. 10.1371/journal.pone.0227841.

36. Stein, K.C., Morales-Polanco, F., van der Lienden, J., Rainbolt, T.K., and Frydman, J. (2022). Ageing exacerbates ribosome pausing to disrupt cotranslational proteostasis. Nature 601, 637–642. 10.1038/s41586-021-04295-4.

37. Escalante, L.E., Hose, J., Howe, H., Paulsen, N., Place, M., and Gasch, A.P. (2024). Premature aging in aneuploid yeast is caused in part by aneuploidy-induced defects in Ribosome Quality Control. bioRxiv. 10.1101/2024.06.22.600216.

38. Martin, P.B., Kigoshi-Tansho, Y., Sher, R.B., Ravenscroft, G., Stauffer, J.E., Kumar, R., Yonashiro, R., Müller, T., Griffith, C., Allen, W., et al. (2020). NEMF mutations that impair ribosome-associated quality control are associated with neuromuscular disease. Nat. Commun. 11, 4625. 10.1038/s41467-020-18327-6.

39. Rimal, S., Li, Y., Vartak, R., Geng, J., Tantray, I., Li, S., Huh, S., Vogel, H., Glabe, C., Grinberg, L.T., et al. (2021). Inefficient quality control of ribosome stalling during APP synthesis generates CAT-tailed species that precipitate hallmarks of Alzheimer’s disease. Acta Neuropathol Commun 9, 169. 10.1186/s40478-021-01268-6.

40. Geng, J., Li, S., Li, Y., Wu, Z., Bhurtel, S., Rimal, S., Khan, D., Ohja, R., Brandman, O., and Lu, B. (2024). Stalled translation by mitochondrial stress upregulates a CNOT4-ZNF598 ribosomal quality control pathway important for tissue homeostasis. Nat. Commun. 15, 1637. 10.1038/s41467-024-45525-3.

41. Sokalingam, S., Raghunathan, G., Soundrarajan, N., and Lee, S.-G. (2012). A study on the effect of surface lysine to arginine mutagenesis on protein stability and structure using green fluorescent protein. PLoS One 7, e40410. 10.1371/journal.pone.0040410.

42. Koren, I., Timms, R.T., Kula, T., Xu, Q., Li, M.Z., and Elledge, S.J. (2018). The Eukaryotic Proteome Is Shaped by E3 Ubiquitin Ligases Targeting C-Terminal Degrons. Cell 173, 1622–1635.e14. 10.1016/j.cell.2018.04.028.

43. Singh, J., Kumar, H., Sabareesan, A.T., and Udgaonkar, J.B. (2014). Rational stabilization of helix 2 of the prion protein prevents its misfolding and oligomerization. J. Am. Chem. Soc. 136, 16704–16707. 10.1021/ja510964t.

44. Abskharon, R., Wang, F., Vander Stel, K.J., Sinniah, K., and Ma, J. (2016). The role of the unusual threonine string in the conversion of prion protein. Sci. Rep. 6, 38877. 10.1038/srep38877.

45. Boyer, D.R., Li, B., Sun, C., Fan, W., Sawaya, M.R., Jiang, L., and Eisenberg, D.S. (2019). Structures of fibrils formed by α-synuclein hereditary disease mutant H50Q reveal new polymorphs. Nat. Struct. Mol. Biol. 26, 1044–1052. 10.1038/s41594-019-0322-y.

46. Izawa, T., Park, S.-H., Zhao, L., Hartl, F.U., and Neupert, W. (2017). Cytosolic Protein Vms1 Links Ribosome Quality Control to Mitochondrial and Cellular Homeostasis. Cell 171, 890–903.e18. 10.1016/j.cell.2017.10.002.

47. Su, T., Izawa, T., Thoms, M., Yamashita, Y., Cheng, J., Berninghausen, O., Hartl, F.U., Inada, T., Neupert, W., and Beckmann, R. (2019). Structure and function of Vms1 and Arb1 in RQC and mitochondrial proteome homeostasis. Nature 570, 538–542. 10.1038/s41586-019-1307-z.

48. Yip, M.C.J., Savickas, S., Gygi, S.P., and Shao, S. (2020). ELAC1 Repairs tRNAs Cleaved during Ribosome-Associated Quality Control. Cell Rep. 30, 2106–2114.e5. 10.1016/j.celrep.2020.01.082.

49. Kelkar, D.A., Khushoo, A., Yang, Z., and Skach, W.R. (2012). Kinetic analysis of ribosome-bound fluorescent proteins reveals an early, stable, cotranslational folding intermediate. J. Biol. Chem. 287, 2568–2578. 10.1074/jbc.M111.318766.

50. Kaiser, C.M., Goldman, D.H., Chodera, J.D., Tinoco, I., Jr, and Bustamante, C. (2011). The ribosome modulates nascent protein folding. Science 334, 1723–1727. 10.1126/science.1209740.

51. Goldman, D.H., Kaiser, C.M., Milin, A., Righini, M., Tinoco, I., and Bustamante, C. (2015). Mechanical force releases nascent chain–mediated ribosome arrest in vitro and in vivo. Science 348, 457–460. 10.1126/science.1261909.

52. Leininger, S.E., Trovato, F., Nissley, D.A., and O’Brien, E.P. (2019). Domain topology, stability, and translation speed determine mechanical force generation on the ribosome. Proc. Natl. Acad. Sci. U. S. A. 116, 5523–5532. 10.1073/pnas.1813003116.

53. Bustamante, C., Alexander, L., Maciuba, K., and Kaiser, C.M. (2020). Single-Molecule Studies of Protein Folding with Optical Tweezers. Annu. Rev. Biochem. 89, 443–470. 10.1146/annurev-biochem-013118-111442.

54. Lu, J., and Deutsch, C. (2008). Electrostatics in the ribosomal tunnel modulate chain elongation rates. J. Mol. Biol. 384, 73–86. 10.1016/j.jmb.2008.08.089.

55. Su, T., Cheng, J., Sohmen, D., Hedman, R., Berninghausen, O., von Heijne, G., Wilson, D.N., and Beckmann, R. (2017). The force-sensing peptide VemP employs extreme compaction and secondary structure formation to induce ribosomal stalling. Elife 6. 10.7554/eLife.25642.

56. Kudva, R., Tian, P., Pardo-Avila, F., Carroni, M., Best, R.B., Bernstein, H.D., and von Heijne, G. (2018). The shape of the bacterial ribosome exit tunnel affects cotranslational protein folding. Elife 7. 10.7554/eLife.36326.

57. Mermans, D., Nicolaus, F., Baygin, A., and von Heijne, G. (2023). Cotranslational folding of human growth hormone in vitro and in Escherichia coli. FEBS Lett. 597, 1355–1362. 10.1002/1873-3468.14562.

58. Chen, X., Rajasekaran, N., Liu, K., and Kaiser, C.M. (2020). Synthesis runs counter to directional folding of a nascent protein domain. Nat. Commun. 11, 5096. 10.1038/s41467-020-18921-8.

59. Bertz, M., and Rief, M. (2008). Mechanical unfoldons as building blocks of maltose-binding protein. J. Mol. Biol. 378, 447–458. 10.1016/j.jmb.2008.02.025.

60. Nilsson, O.B., Nickson, A.A., Hollins, J.J., Wickles, S., Steward, A., Beckmann, R., von Heijne, G., and Clarke, J. (2017). Cotranslational folding of spectrin domains via partially structured states. Nat. Struct. Mol. Biol. 24, 221–225. 10.1038/nsmb.3355.

61. Kemp, G., Nilsson, O.B., Tian, P., Best, R.B., and von Heijne, G. (2020). Cotranslational folding cooperativity of contiguous domains of α-spectrin. Proc. Natl. Acad. Sci. U. S. A. 117, 14119–14126. 10.1073/pnas.1909683117.

62. Jaud, S., Fernández-Vidal, M., Nilsson, I., Meindl-Beinker, N.M., Hübner, N.C., Tobias, D.J., von Heijne, G., and White, S.H. (2009). Insertion of short transmembrane helices by the Sec61 translocon. Proc. Natl. Acad. Sci. U. S. A. 106, 11588–11593. 10.1073/pnas.0900638106.

63. Ismail, N., Hedman, R., Schiller, N., and von Heijne, G. (2012). A biphasic pulling force acts on transmembrane helices during translocon-mediated membrane integration. Nat. Struct. Mol. Biol. 19, 1018–1022. 10.1038/nsmb.2376.

64. Ismail, N., Hedman, R., Lindén, M., and von Heijne, G. (2015). Charge-driven dynamics of nascent-chain movement through the SecYEG translocon. Nat. Struct. Mol. Biol. 22, 145–149. 10.1038/nsmb.2940.

65. Kriegler, T., Magoulopoulou, A., Amate Marchal, R., and Hessa, T. (2018). Measuring Endoplasmic Reticulum Signal Sequences Translocation Efficiency Using the Xbp1 Arrest Peptide. Cell Chem Biol 25, 880–890.e3. 10.1016/j.chembiol.2018.04.006.

66. Verma, R., Oania, R.S., Kolawa, N.J., and Deshaies, R.J. (2013). Cdc48/p97 promotes degradation of aberrant nascent polypeptides bound to the ribosome. Elife 2, e00308. 10.7554/eLife.00308.

67. Sá-Moura, B., Simões, A.M., Fraga, J., Fernandes, H., Abreu, I.A., Botelho, H.M., Gomes, C.M., Marques, A.J., Dohmen, R.J., Ramos, P.C., et al. (2013). Biochemical and biophysical characterization of recombinant yeast proteasome maturation factor ump1. Comput. Struct. Biotechnol. J. 7, e201304006. 10.5936/csbj.201304006.

68. Fritch, B., Kosolapov, A., Hudson, P., Nissley, D.A., Woodcock, H.L., Deutsch, C., and O’Brien, E.P. (2018). Origins of the Mechanochemical Coupling of Peptide Bond Formation to Protein Synthesis. J. Am. Chem. Soc. 140, 5077–5087. 10.1021/jacs.7b11044.

69. Leininger, S.E., Rodriguez, J., Vu, Q.V., Jiang, Y., Li, M.S., Deutsch, C., and O’Brien, E.P. (2021). Ribosome Elongation Kinetics of Consecutively Charged Residues Are Coupled to Electrostatic Force. Biochemistry 60, 3223–3235. 10.1021/acs.biochem.1c00507.

70. Joiret, M., Kerff, F., Rapino, F., Close, P., and Geris, L. (2022). Ribosome exit tunnel electrostatics. Phys Rev E 105, 014409. 10.1103/PhysRevE.105.014409.

71. Fedyukina, D.V., and Cavagnero, S. (2011). Protein folding at the exit tunnel. Annu. Rev. Biophys. 40, 337–359. 10.1146/annurev-biophys-042910-155338.

72. Dao Duc, K., Batra, S.S., Bhattacharya, N., Cate, J.H.D., and Song, Y.S. (2019). Differences in the path to exit the ribosome across the three domains of life. Nucleic Acids Res. 47, 4198–4210. 10.1093/nar/gkz106.

73. Lu, J., and Deutsch, C. (2005). Folding zones inside the ribosomal exit tunnel. Nat. Struct. Mol. Biol. 12, 1123–1129. 10.1038/nsmb1021.

74. Blaber, M., Zhang, X.J., and Matthews, B.W. (1993). Structural basis of amino acid alpha helix propensity. Science 260, 1637–1640. 10.1126/science.8503008.

75. Chakrabartty, A., and Baldwin, R.L. (1995). Stability of alpha-helices. Adv. Protein Chem. 46, 141–176.

76. Pace, C.N., and Scholtz, J.M. (1998). A helix propensity scale based on experimental studies of peptides and proteins. Biophys. J. 75, 422–427. 10.1016/s0006-3495(98)77529-0.

77. Meller, A., Coombs, D., and Shalgi, R. (2020). Fluorescent polysome profiling reveals stress-mediated regulation of HSPA14-ribosome interactions. bioRxiv, 860833. 10.1101/860833.

78. Martin, T.E., and Hartwell, L.H. (1970). Resistance of active yeast ribosomes to dissociation by KCl. J. Biol. Chem. 245, 1504–1506.

79. Gamerdinger, M., Kobayashi, K., Wallisch, A., Kreft, S.G., Sailer, C., Schlömer, R., Sachs, N., Jomaa, A., Stengel, F., Ban, N., et al. (2019). Early Scanning of Nascent Polypeptides inside the Ribosomal Tunnel by NAC. Mol. Cell 75, 996–1006.e8. 10.1016/j.molcel.2019.06.030.

80. Bhushan, S., Hoffmann, T., Seidelt, B., Frauenfeld, J., Mielke, T., Berninghausen, O., Wilson, D.N., and Beckmann, R. (2011). SecM-stalled ribosomes adopt an altered geometry at the peptidyl transferase center. PLoS Biol. 9, e1000581. 10.1371/journal.pbio.1000581.

81. Huter, P., Arenz, S., Bock, L.V., Graf, M., Frister, J.O., Heuer, A., Peil, L., Starosta, A.L., Wohlgemuth, I., Peske, F., et al. (2017). Structural Basis for Polyproline-Mediated Ribosome Stalling and Rescue by the Translation Elongation Factor EF-P. Mol. Cell 68, 515–527.e6. 10.1016/j.molcel.2017.10.014.

82. Jiang, Y., and O’Brien, E.P. (2021). Mechanical Forces Have a Range of Effects on the Rate of Ribosome Catalyzed Peptidyl Transfer Depending on Direction. J. Phys. Chem. B 125, 7128–7136. 10.1021/acs.jpcb.1c02263.

83. Wilson, D.N., Arenz, S., and Beckmann, R. (2016). Translation regulation via nascent polypeptide-mediated ribosome stalling. Curr. Opin. Struct. Biol. 37, 123–133. 10.1016/j.sbi.2016.01.008.

84. Bhunia, S., Singh, A., and Ojha, A.K. (2016). Un-catalyzed peptide bond formation between two monomers of glycine, alanine, serine, threonine, and aspartic acid in gas phase: a density functional theory study. Eur. Phys. J. D 70, 106. 10.1140/epjd/e2016-60609-8.

85. Hentzen, D., Mandel, P., and Garel, J.P. (1972). Relation between aminoacyl-tRNA stability and the fixed amino acid. Biochim. Biophys. Acta 281, 228–232. 10.1016/0005-2787(72)90174-8.

86. Spirin, A.S. (1985). Ribosomal translocation: facts and models. Prog. Nucleic Acid Res. Mol. Biol. 32, 75–114. 10.1016/s0079-6603(08)60346-3.

87. Noller, H. (2006). 10 evolution of ribosomes and translation from an RNA World. Cold Spring Harbor Monograph Archive 43, 287–307. 10.1101/087969739.43.287.

88. Ramakrishnan, V. (2009). The ribosome: some hard facts about its structure and hot air about its evolution. Cold Spring Harb. Symp. Quant. Biol. 74, 25–33. 10.1101/sqb.2009.74.032.

89. Trentini, D.B., Pecoraro, M., Tiwary, S., Cox, J., Mann, M., Hipp, M.S., and Hartl, F.U. (2020). Role for ribosome-associated quality control in sampling proteins for MHC class I-mediated antigen presentation. Proc. Natl. Acad. Sci. U. S. A. 117, 4099–4108. 10.1073/pnas.1914401117.

90. Schmidt, C., Becker, T., Heuer, A., Braunger, K., Shanmuganathan, V., Pech, M., Berninghausen, O., Wilson, D.N., and Beckmann, R. (2016). Structure of the hypusinylated eukaryotic translation factor eIF-5A bound to the ribosome. Nucleic Acids Res. 44, 1944–1951. 10.1093/nar/gkv1517.

91. Petukhov, M., Uegaki, K., Yumoto, N., Yoshikawa, S., and Serrano, L. (1999). Position dependence of amino acid intrinsic helical propensities II: non-charged polar residues: Ser, Thr, Asn, and Gln. Protein Sci. 8, 2144–2150. 10.1110/ps.8.10.2144.

